# Dual Translational Control in Cardiomyocytes by Heterogeneous mTORC1 and Hypertrophic ERK Activation

**DOI:** 10.1101/2025.02.10.635974

**Authors:** Keita Uchida, Emily A. Scarborough, Benjamin L. Prosser

**Affiliations:** Department of Physiology, Pennsylvania Muscle Institute, University of Pennsylvania Perelman School of Medicine, Philadelphia, PA 19104, USA

**Keywords:** single-cell protein synthesis, cardiac hypertrophy, translation initiation, ribosome biogenesis, 4E-BP1

## Abstract

**Background:** Cardiac hypertrophy allows post-mitotic cardiomyocytes to meet increased hemodynamic demands but can predispose the heart to adverse clinical outcomes. Despite its central role in cardiac adaptation, the translational control mechanisms that drive cardiac hypertrophy are poorly understood. In this study, we elucidate the relative contributions of the various translational control mechanisms operant during homeostasis and hypertrophic growth.

**Methods:** A combination of immunofluorescence and single myocyte protein synthesis assays were used to dissect the single-cardiomyocyte mechanisms of translational control under basal and hypertrophic conditions in isolated adult rat cardiomyocytes. Translational control mechanism were examined in a mouse model of acute hypertrophic phenylephrine (PE) stimulation prior to overt cardiac growth.

**Results:** We observed strikingly heterogeneous activity of mTORC1, the master regulator of translation, across cardiomyocytes both *in situ* and *ex vivo*. Heterogeneous mTORC1 activity drove heterogeneous protein synthesis, with translation primarily controlled via canonical mTORC1-dependent 4EBP1 phosphorylation at Thr36/Thr45/Thr69 under baseline conditions. Hypertrophic PE stimulation recruited more cardiomyocytes into a high mTORC1 activity state. PE induced a switch in 4EBP1 phosphorylation by increasing mTORC1-dependent phosphorylation at Thr36/Thr45, but not Thr69. Further, PE induced a novel mTORC1-independent, but MEK-ERK-dependent, pathway driving 4EBP1 phosphorylation at Ser64 in both isolated cardiomyocytes and *in vivo*. Ribosome biogenesis was also observed within hours upon hypertrophic stimulation, while the mTORC1-S6K-eEF2K-eEF2 pathway was not found to be a major driver of protein translation.

**Conclusions:** Protein synthesis is heterogeneous across cardiomyocytes driven by heterogeneous mTORC1 activity. MEK-ERK signaling directly controls 4EBP1 phosphorylation to augment translation during cardiac hypertrophy and challenges the canonical model of translation initiation.

**Key points:**

- mTORC1 activity is low and heterogeneous across cardiomyocytes at baseline.
- Heterogeneous mTORC1 activity drives variable 4EBP1 phosphorylation at Thr36/Thr45/Thr69, resulting in heterogeneous protein translation.
- Phenylephrine stimulation recruits more cardiomyocytes into a high mTORC1 activity state to augment protein synthesis through 4EBP1 phosphorylation at Thr36/Thr45, but not Thr69.
- Phenylephrine stimulation boosts protein translation through mTORC1-independent but ERK-dependent phosphorylation of 4EBP1 at Ser64.
- mTORC1-S6K-eEF2K-eEF2 pathway is not a major driver of global protein translation.
- Ribosome biogenesis is observed within hours after hypertrophic stimulation.
- The data demonstrates a dual input model of mTORC1 and ERK dependent phosphorylation of 4EBP1 to regulate protein translation in cardiomyocytes.

## INTRODUCTION

Cell size and growth rate are determined by the rates of accumulation and loss of macromolecules ^1^. The majority of work studying cell size has been performed in proliferating cell lines, which primarily use their energy to generate new proteins to grow ^2^. In contrast, cardiomyocytes are long-lived, terminally differentiated cells that must balance the energetic demands of constant contraction with macromolecule synthesis. The heart displays some of the lowest protein synthesis rates of any organ in the body ^3^ but must rapidly activate its protein synthesis machinery to adapt to increases in hemodynamic load and stress. However, excessive cardiac growth (hypertrophy) predisposes the heart to arrhythmias, contractile dysfunction, and ultimately failure, and thus understanding cardiomyocyte size regulation is critical for human health and longevity. Despite these unique features of cardiomyocyte size regulation, the translational control mechanisms underlying homeostasis and dynamic activation of protein synthesis during hypertrophic stimulation remain incompletely understood.

In the heart, cardiomyocytes are exposed to mechanical and neurohormonal cues that regulate their growth. Diverse hypertrophic stimuli activate distinct signaling pathways that ultimately converge on mTORC1^4–6^, a key kinase that integrates amino acid and energy availability with extracellular signals to regulate protein synthesis (Supplemental Figure 1) ^7,8^. In response to a stressor, the heart undergoes two phases of translational control mechanisms to promote hypertrophy ^9^. First, there is an acute increase in *translational efficiency* by promotion of translation initiation and elongation. mTORC1 phosphorylation of 4EBP1, a negative repressor of translation initiation, leads to its dissociation from the 5’ cap of mRNA thereby allowing for the formation of the eIF4F translation initiation complex (Figure 1A) ^10–12^. Additionally, mTORC1 activates S6K, which then phosphorylates eEF2K leading to its inactivation and subsequent dephosphorylation of eEF2 ^13,14^. Dephosphorylated eEF2 is associated with an accelerated translation elongation rate ^15^. These mechanisms accelerate the loading of ribosomes onto mRNA and the transit rate of ribosomes across the coding sequence, respectively. This acute acceleration of translation rate is followed by a secondary increase in *translational capacity* to maintain an increased rate of protein synthesis. mTORC1 activation is associated with preferential translation of 5’ TOP motif containing mRNA, particularly those encoding ribosomal proteins, small nuclear ribonucleoproteins, and translation factors, to drive ribosome biogenesis and increase the abundance of translational machinery ^16^. While these mechanisms are well characterized independently, the relative contribution and coordination of these downstream effectors in the heart remain unclear.

**Figure 1:**
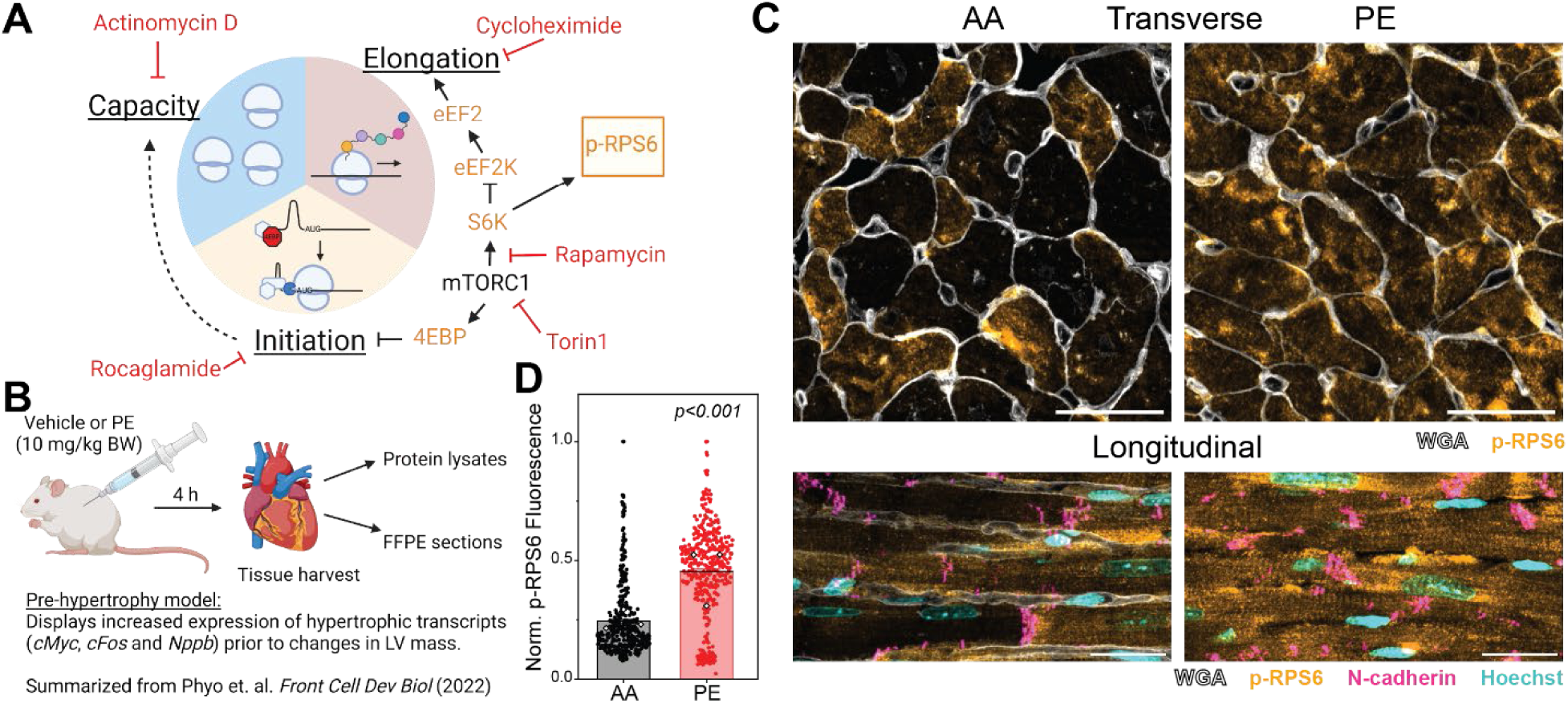
mTORC1 activity in the heart is heterogeneous under basal conditions and is activated by PE injection. (A) Diagram of translational control mechanisms regulated by mTORC1 activity. mTORC1 activity can be measured by monitoring downstream phosphorylation of RPS6 (pRPS6). Pharmacological inhibitors of specific pathways used in later figures are marked in red. (B) Schematic depicting the pre-hypertrophy model. This acute PE injection model ^22^ activates expression of hypertrophic transcripts before cardiac growth is detected. (C) Representative myocardial sections from mouse hearts isolated after AA (left) or PE (right) treatment. Regions with transversally (top) and longitudinally (bottom) oriented cardiomyocytes are shown. Cell borders are labeled with WGA (white) and mTORC1 activity with pRPS6 (orange). In longitudinal sections, intercalated disks marked by N-cadherin (magenta) and nuclei by Hoechst (cyan) are also shown. Scale bars represent 20 µm. (D) Quantification of cellular pRPS6 fluorescence from transverse sections normalized to the maximum cellular fluorescence intensity of each animal to emphasize the distribution shift with PE treatment. Individual cell measurements are displayed as closed circles, mean values from each biological replicate are shown in open diamonds. Mean + standard deviation; (N = 3 hearts). Statistical significance determined using a two-sample t-test with Welch correction (equal variance is not assumed).

Downstream 4EBP1 phosphorylation has not been systematically examined in cardiomyocytes and is largely assumed from studies in other cell types. When unphosphorylated or hypophosphorylated, 4EBP1 sequesters eIF4E to prevent translation initiation. Variable mTORC1 activity fine tunes protein synthesis rates by phosphorylating 4EBP1 at multiple residues ^7^ (Supplemental Figure 1). Canonical 4EBP1 phosphorylation is thought to proceed hierarchically^17^: phosphorylation at “priming sites” Thr36 and Thr45 is required for further phosphorylation at mitogen sensitive sites, Thr69 followed by Ser64 (numbering for rodents will be used, add 1 to each residue number for human 4EBP) ^17^. Mechanistically, phosphorylation of 4EBP1 at the priming sites induces a conformational change that partially disrupts eIF4E binding ^18^ and also positions the distal sites near the mTORC1 kinase active site ^19^. Subsequent phosphorylation of any one of the distal sites (Thr69, Ser64, or Ser82) fully frees eIF4E from 4EBP1 binding ^20^, and allows for the recruitment of other translation initiation factors culminating in the loading of ribosomes to mRNA. Currently, this mTORC1 dependent 4EBP1 phosphorylation model is the sole model assumed operant in cardiomyocytes and alternative pathways have not been explored. Further, protein synthesis and mTORC1 regulation have largely been studied in aggregate using biochemical approaches that lack single-cell resolution.

Here, we sought to examine the single cell mechanisms of translational control at baseline and upon hypertrophic activation. We observed surprisingly heterogenous mTORC1 activity between cardiomyocytes *in situ* under baseline conditions. This considerable variation in mTORC1 activity resulted in heterogeneous protein translation in isolated cardiomyocytes. The heterogeneity in protein synthesis rate at baseline was driven primarily by mTORC1 dependent phosphorylation of 4EBP1 at Thr36/45/69. Hypertrophic stimulation with PE increased mTORC1 activity both in isolated cardiomyocytes and *in vivo,* resulting in protein synthesis activation through three key mechanisms: (1) canonical mTORC1-dependent phosphorylation of 4EBP1 at Thr36 and Thr45, but not Thr69. (2) novel mTORC1-independent but ERK-dependent phosphorylation of 4EBP1 Ser64 and (3) rapid activation of ribosome biogenesis. Inhibition of the S6K pathway had little to no observable effect on protein synthesis, suggesting a minor role for elongation rate in regulating global translation. Overall, the results define the relative contributions of translational control mechanisms during homeostasis and hypertrophic growth and present a novel, dual input model of 4EBP1 phosphorylation to control translation rate.

## MATERIALS AND METHODS

### Animals

Animal care and use procedures were performed in accordance with the standards set forth by the University of Pennsylvania Institutional Animal Care and Use Committee and the Guide for the Care and Use of Laboratory Animals published by the US National Institutes of Health; protocols were approved by the University of Pennsylvania Institutional Animal Care and Use Committee.

### Adult rat cardiomyocyte isolation and culture

Primary adult ventricular myocytes were isolated from 8- to 12-week-old Sprague Dawley rats using Langendorff retrograde aortic perfusion with an enzymatic solution as previously described ^21^. Briefly, the heart was removed from an anesthetized rodent under isoflurane and retrograde-perfused on a Langendorff apparatus with a collagenase solution. The digested heart was then minced and triturated with glass pipettes to free individual cardiomyocytes. The resulting supernatant was separated and centrifuged at 300 rpm to isolate cardiomyocytes. These cardiomyocytes were then resuspended in aRVM media (Medium 199 (Thermo Fisher) supplemented with 1x insulin-transferrin-selenium-X (ITS, Gibco), 1 μg/μL primocin (InvivoGen), and 20 mM HEPES, pH = 7.4 (UPenn Cell Center)) at low density, cultured at 37 °C and 5% CO2 with the addition of 25 μM of cytochalasin D (Cayman Chemical) in the media.

### Preparation of lysates from 4h AA or PE treated mice

Male and female C57BL/6 mice (9-10 weeks old) were subcutaneously injected with ascorbic acid vehicle (AA) or PE (10 mg/kg BW) around 9-11 AM as previously described ^22^. Randomized pairs of animals were injected with vehicle or PE in an alternating manner and were staggered to ensure that the hearts were extracted 4 hours after the injection time. Mice were anesthetized with 3% isoflurane, heparinized, and hearts were rapidly extracted and placed in ice cold PBS. The atria were removed, and small 20-30 mg pieces of ventricular tissue were weighed and flash frozen in cryotubes. Lysates were prepared by homogenizing the tissue pieces in 50 µL/mg of 1X RIPA + 2% SDS + 1x protease/phosphatase inhibitors + EDTA (1 mM). Protein concentration was measured by BCA assay (Thermo Scientific Pierce). Supernatant was mixed with 4X sample buffer (LICOR) with β-mercaptoethanol and warmed at 37°C for 30 minutes. No hearts/animals were excluded from the study and blinding was not performed.

### Preparation of mouse hearts for immunohistochemistry and measurement of mTORC1 activity

Hearts from C57BL/6 mice treated with AA or PE for 4 hours were removed and perfused with 1X PBS briefly to remove residual blood using Langendorff retrograde aortic perfusion. The solution line was switched to 4% paraformaldehyde (PFA) in PBS and the hearts were perfusion fixed for 15 minutes before the atria were removed and stored in 4% PFA overnight at 4 °C. The following day, the hearts were sequentially washed in PBS, 50% ethanol, and 70% ethanol and stored at 4 °C until processing. The hearts were then processed and embedded in paraffin at the Penn Center for Musculoskeletal Disorders histology core.

Transverse sections of 7 µm thickness across the ventricles were cut. The sections were then processed as described previously^23^. Briefly, the sections were deparaffinized in a series of xylene and ethanol washes. Antigen retrieval was performed with the Reveal Decloaker (Biocare Medical) in a pressure cooker on high heat for 15 minutes. The tissue sections were permeabilized in 0.25% Triton X-100 in PBS, blocked in 3% BSA in PBS + 0.1% Tween-20 (PBST), and incubated with mouse anti-N-Cadherin (Clone 5D5, Abcam ab98952, 1:100) and rabbit anti-phospho-RPS6 (Cell Signaling Technology #5364S, 1:50) in blocking solution overnight at room temperature in a humidity chamber. The samples were washed with PBST and stained with goat anti-rabbit Alexa Fluor 647 (Thermo Fisher Scientific, 1:100), goat anti-mouse Alexa Fluor 568 (Thermo Fisher Scientific, 1:100), and Alexa Fluor 488 conjugated WGA (Thermo Fisher Scientific, 1:400). Sections were stained with Hoechst 33342, trihydrochloride (Invitrogen, 1:1000) before mounting with ProLong Diamond antifade mountant (Invitrogen).

Four channel, 4 x 1 um z-stack images were obtained on a Zeiss 880 Airyscan microscope using a 63x 1.4NA oil immersion objective in fast mode. Maximum intensity projections of the WGA channel were thresholded to generate a binary mask of cell outlines from transverse sections. Particle analysis was performed on the cell outlines to generate single cardiomyocyte ROIs used to measure pRPS6 fluorescence.

### Preparation of lysates from aRVMs

Treated aRVMs were collected in a 1.5 mL microcentrifuge tube, pelleted with gentle tabletop microcentrifugation, and lysed in 1X RIPA + 2% SDS + 1x protease/phosphatase inhibitors. Protein concentration was measured using Pierce BCA protein assay (Thermo Fisher Scientific). Supernatant was mixed with 4X sample buffer (LICOR) with β- mercaptoethanol and boiled at 97°C for 10 minutes.

### Western blotting

Lysates were loaded in equal amounts in each lane and separated on a 4-15% gradient, 15-well, Mini-PROTEAN TGX gel (Bio-Rad). Proteins were transferred to a nitrocellulose membrane using the turbo transfer system (Bio-Rad). Membranes were rinsed in water, blocked in Intercept (TBS) blocking buffer (LI-COR Biosciences), and incubated in primary antibodies (see table below for concentrations) overnight at 4°C. Membranes were washed 3x with TBST (Cell Signaling Technology, #9997S) and incubated with secondary antibodies for 2-4 hours. Membranes were washed 3x with TBST and imaged on an Odyssey Fc (LI-COR). Bands were quantified using the Image Studio software (LI-COR).

In some cases, the membranes were cut horizontally to blot for multiple antibodies targeting different sized proteins but all bands were normalized to the GAPDH loading control from the same membrane. For phospho-protein quantification, the phospho-specific antibody and the total substrate protein were often run separately on different membrane and the ratio of the GAPDH normalized band intensities to the GAPDH normalized band intensities of the total substrate protein are presented. Male and female samples were run on separate gels and the quantification was normalized to the vehicle control values of each group.

### Puromycin incorporation in isolated aRVMs

Stock solutions (1000X) of rapamycin (Sigma-Aldrich. R0395), torin1 (Sigma-Aldrich #475991), U0126 (Sigma-Aldrich 19-147), SCH77284 (ApexBio #B5866), rocaglamide (Cayman Chemical #14841), cycloheximide (Sigma-Aldrich 01810), LY294002 (Santa Cruz Biotechnology sc-201426), and Actinomycin D (Sigma-Aldrich A9415) were prepared in DMSO and aliquots were stored at −20 °C until use. Phenylephrine and isoproterenol (Sigma-Aldrich #I6504) were prepared in media immediately before use. Isolated aRVMs were attached to MyoTak (IonOptix) coated glass coverslips in 12-well tissue culture plates. To insulin starve aRVMs, cells were washed twice in M199 media and incubated in ITS- free aRVM media overnight (∼18 hours). Cardiomyocytes were treated with inhibitors for 15 minutes prior to stimulation with phenylephrine (Sigma-Aldrich P6126, lot# SLCH3274, 10 µM) for 2 hours. Afterwards, cardiomyocytes were washed with pre-warmed media containing the treatment and puromycin (Sigma-Aldrich P8833, 1 µM) for 15 minutes. The solution exchanges were performed rapidly using a vacuum and all treatments were pre-warmed to 37°C to minimize sudden temperature changes. Afterwards, aRVMs were washed once in PBS, fixed with 4% paraformaldehyde, permeabilized in 0.25% triton X-100 in PBS, and blocked in 3% BSA (Sigma Aldrich) in PBS. Samples were incubated in primary antibodies (See Antibodies and concentrations for IF in aRVMs below) overnight at RT. Samples were washed in PBS and incubated in secondary antibodies (1:500) overnight at RT. Samples were stained with Hoechst 33342,trihydrochloride (Invitrogen, 1:1000) before mounting with Prolong diamond antifade mountant (Invitrogen).

### HPG incorporation in isolated aRVMs

aRVMs were plated onto MyoTak (Ionoptix) coated coverslips placed in 12-well plates and cultured overnight in aRVM media. The following morning, the media was washed twice in methionine free RPMI media (Thermo Fisher Scientific) and aRVMs were cultured for 2 hours in complete methionine free RPMI media containing HEPES, primocin, and cytochalasin D with the indicated pharmacological treatments. After the methionine depletion step, the media was exchanged with complete methionine free media containing HPG (Click Chemistry Tools, 100 µM) and with the indicated pharmacological treatments. After the indicated HPG incorporation time, aRVMs were washed once in PBS and fixed in 4% paraformaldehyde (Electron Microscopy Sciences) for 15 min., then permeabilized in PBS with 0.25% Triton X-100. HPG labeling was performed using the Click-&-Go Plus 488 imaging kit (Click Chemistry Tools) according to the manufacturer instructions.

### Widefield microscopy

Tile scan images of aRVMs stained for puromycin or HPG and pRPS6 were obtained on a Zeiss AxioObserver Microscope using a 20X air 0.8 NA objective (Cell and Developmental Biology Microscopy Core). Z-stacks consisting of 5 optical slices at 2 um intervals (8 um total) were imaged across 24 tiles (4x6), resulting in an ∼ 2 mm x 2 mm image. The focal plane was adjusted at 16 support points (4x4 grid) throughout the tile scan to correct for potential focal drift. No binning was performed and a 1x digital zoom was used, resulting in pixel size of 293 nm x 293 nm. Samples were sequentially excited with LED light at 567nm and 630nm and emission filters collected light between 580-610nm and 663-733nm, respectively. Images were stitched together using the Zen software (Carl Zeiss AG).

### Tile scan imaging of on Zeiss 980

A 20X 0.8 NA air objective was used to image fixed coverslips of aRVMs stained for puromycin and phosphosite specific 4EBP1 antibodies. Z-stacks consisting of 4 optical slices at 2 um intervals (6 um total) were imaged across 25 tiles (5x5), resulting in an image ∼ 1.15 mm x 1.15 mm. The focal plane was adjusted at 16 support points (4x4 grid) throughout the tile scan to correct for potential focal drift. A digital zoom of 1.7x was used resulting in a pixel size of 62 x 62 nm. The images were acquired on the 4Y Fast Airyscan mode. Each channel was imaged sequentially each z-stack with the following excitation wavelength and emission filters to minimize fluorescence crosstalk: 405 nm/ short pass 505 nm; 488 nm / short pass 550 nm, 561 nm / short pass 615 nm, and 639 nm/ long pass 570 nm. Images were Airyscan processed and then stitched together using the Zen software (Carl Zeiss AG).

### Quantification of widefield and tile scan images

Maximum intensity projections were prepared from the tile scan images. Preliminary regions of interests (ROIs) were obtained by segmenting tile scan images of aRVMs with automated thresholding in FIJI. The identified ROIs were quality controlled by separating fused ROIs from multiple aRVMs, excluding ROIs originating from dead cells or debris, and removing ROIs from overlapping aRVMs. The mean fluorescence intensities from the remaining ROIs were measured from the maximum intensity projections. To quantify normalized puromycin incorporation and phospho-RPS6, aRVMs treated with secondary antibodies only were prepared for background subtraction. The measured cellular fluorescence was normalized to the average fluorescence from the DMSO group. Normalization and quantification were performed using the OriginPro 2019 software (OriginLab).

### Quantification of methionine abundance in mouse and human proteomes

Manually curated human and mouse proteomes were downloaded from UniProt Knowledgebase/Swiss-Prot (www.uniprot.org) on 8/18/2024. A total of 20,435 and 17,217 proteins from the human and mouse proteomes, respectively, were analyzed in MATLAB (MathWorks). The total abundances of each amino acid were counted across the proteomes and normalized to the sum of amino acid count in Supplemental Figure 2A. The number of methionines per protein were counted and the raw counts (left) or counts normalized to the protein length (right) are displayed as histograms in Supplemental Figure 2B.

### Antibodies and concentrations for Western Blotting

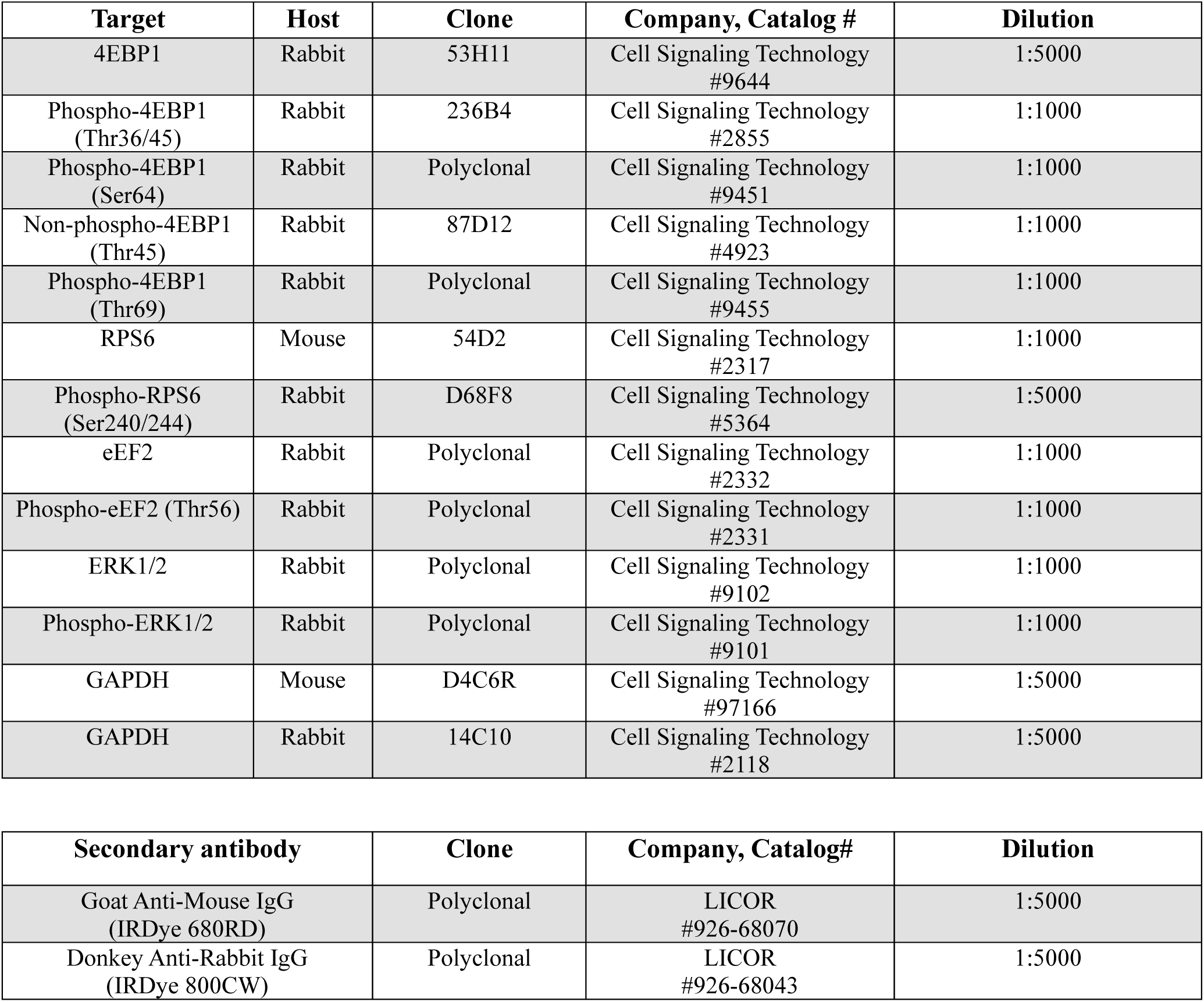

**Antibodies and concentrations for Western Blotting:**
| **Target** | **Host** | **Clone** | **Company, Catalog #** | **Dilution** |
| --- | --- | --- | --- | --- |
| 4EBP1 | Rabbit | 53H11 | Cell Signaling Technology<br>#9644 | 1:5000 |
| Phospho-4EBP1<br>(Thr36/45) | Rabbit | 236B4 | Cell Signaling Technology<br>#2855 | 1:1000 |
| Phospho-4EBP1<br>(Ser64) | Rabbit | Polyclonal | Cell Signaling Technology<br>#9451 | 1:1000 |
| Non-phospho-4EBP1<br>(Thr45) | Rabbit | 87D12 | Cell Signaling Technology<br>#4923 | 1:1000 |
| Phospho-4EBP1<br>(Thr69) | Rabbit | Polyclonal | Cell Signaling Technology<br>#9455 | 1:1000 |
| RPS6 | Mouse | 54D2 | Cell Signaling Technology<br>#2317 | 1:1000 |
| Phospho-RPS6<br>(Ser240/244) | Rabbit | D68F8 | Cell Signaling Technology<br>#5364 | 1:5000 |
| eEF2 | Rabbit | Polyclonal | Cell Signaling Technology<br>#2332 | 1:1000 |
| Phospho-eEF2 (Thr56) | Rabbit | Polyclonal | Cell Signaling Technology<br>#2331 | 1:1000 |
| ERK1/2 | Rabbit | Polyclonal | Cell Signaling Technology<br>#9102 | 1:1000 |
| Phospho-ERK1/2 | Rabbit | Polyclonal | Cell Signaling Technology<br>#9101 | 1:1000 |
| GAPDH | Mouse | D4C6R | Cell Signaling Technology<br>#97166 | 1:5000 |
| GAPDH | Rabbit | 14C10 | Cell Signaling Technology<br>#2118 | 1:5000 |

| **Secondary antibody** | **Clone** | **Company, Catalog#** | **Dilution** |
| --- | --- | --- | --- |
| Goat Anti-Mouse IgG<br>(IRDye 680RD) | Polyclonal | LICOR<br>#926-68070 | 1:5000 |
| Donkey Anti-Rabbit IgG<br>(IRDye 800CW) | Polyclonal | LICOR<br>#926-68043 | 1:5000 |

### Antibodies and concentrations for IF in aRVMs

**Antibodies and concentrations for IF in aRVMs:**
| **Target** | **Host** | **Clone** | **Company, Catalog #** | **Dilution** |
| --- | --- | --- | --- | --- |
| Puromycin | Mouse | 12D10 | EMD Millipore #MABE343 | 1:250 |
| Phospho-RPS6<br>(Ser 240/244) | Rabbit | D68F8 | Cell Signaling Technology #5364S | 1:250 |
| Phospho-mTOR<br>(Ser 2448) | Rabbit | Poly | Cell Signaling Technology #2971S | 1:250 |
| p4EBP1 (Thr36/45) | Rabbit | 236B4 | Cell Signaling Technology #2855 | 1:250 |
| Nonphospho-4EBP1 (Thr46) | Rabbit | 87D12 | Cell Signaling Technology #4923 | 1:250 |
| Phospho-4EBP1 (Thr69) | Rabbit | Polyclonal | Cell Signaling Technology #9455 | 1:250 |

## RESULTS

### mTORC1 activity is low and heterogeneous and augmented by hypertrophic stimulation *in situ*

We previously described a reliable model of cardiac hypertrophy induced by three injections of PE over 5 days ^21^. After the final injection, the heart weight to tibia length increases by ∼ 30% with concomitant increase in protein synthesis rate. When the hearts are examined just 4 hours after the initial injection, markers of hypertrophy induction are observed including robust increases in the expression of hypertrophic genes *cMyc*, *cFos*, and *Nppb* but before the detection of cardiac growth ^22^, demonstrating its utility as a model to study the signaling pathways leading to hypertrophic remodeling (Figure 1B). We began by examining mTORC1 activity in tissue sections collected from animals 4 hours after a single injection of PE or vehicle. Mouse left ventricular tissue sections were stained for phosphorylated ribosomal protein S6 (pRPS6), an established marker of mTORC1 activity. Surprisingly, in transverse cardiomyocyte sections from both endo- and epicardial regions, a mosaic pattern of mTORC1 activation was observed with many cells displaying near background levels of pRPS6 staining and others brightly labeled (Figure 1C). In mid-myocardial regions where cardiomyocytes are longitudinally oriented, individual cardiomyocytes demonstrated uniformly low pRPS6 intensity while immediately adjacent cells exhibited high pRPS6 fluorescence (and no clear gradient in fluorescence, excluding the possibility of technical variability in antibody labeling). In tissue sections from animals treated with PE, the pRPS6 intensity displays a more uniform labeling. Quantification of the normalized cellular fluorescence reveals a bi-modal distribution of pRPS6 with a shift in the population distribution to more cells displaying high pRPS6 intensity in hearts of animals treated with PE (Figure 1D). Overall, the data shows that mTORC1 activity is low and heterogeneous under basal conditions but robustly increased following hypertrophic stimulation with PE *in vivo*.

### mTORC1 activity and protein synthesis are heterogeneous and strongly correlated in isolated cardiomyocytes

The *in situ* observation of heterogeneous mTORC1 activity increased by hypertrophic stimulation raised two key questions: 1) Does heterogeneous mTORC1 activity control differential protein synthesis across cardiomyocytes, and by which downstream mechanism(s)? 2) During early hypertrophic stimulation, what are the relative contributions of different translational control mechanisms to increased protein synthesis?

To determine if the mTORC1 heterogeneity observed *in vivo* is preserved *ex vivo* and reflects heterogeneous protein synthesis rates, we began by measuring single cell protein synthesis rate in isolated, insulin-starved, adult rat ventricular cardiomyocytes (aRVMs) by pulse labeling with puromycin, a tyrosyl-tRNA analog that is covalently attached to growing polypeptides (Figure 2A) ^24^. Following fixation, aRVMs were labeled with antibodies against the puromycylated peptide and pRPS6 to measure both protein synthesis and mTORC1 activity, respectively. Indeed, individual cardiomyocytes displayed a wide range of puromycin incorporation from 8% to 170% of the mean and similarly variable pRPS6 fluorescence (range: 34% to 152% of mean, Figure 2B and C) at baseline. The observed puromycin fluorescence is specific to reporting global translation rates since pretreatment with the translation elongation inhibitor cycloheximide (CHX) ^25^ reduced puromycin fluorescence nearly to background levels (Figure 2C). Similarly, pretreatment of aRVMs with Rocaglamide A (RocA), a small molecule that selectively inhibits cap-dependent translation initiation ^26^, recapitulated the CHX results. Importantly, RocA treatment does not inhibit scanning-independent IRES mediated translation ^26^, indicating that the observed puromycin fluorescence reflects cap-dependent translation. Stimulation of aRVMs with PE for 2 hours shifted the distribution of pRPS6 upwards by 22%, demonstrating the recruitment of more cells into a high mTORC1 activity state. PE also increased puromycin incorporation by 18%, consistent with past studies utilizing radiolabeled amino acids ^27,28^. The heterogeneous pRPS6 fluorescence was strongly correlated with puromycin labeling (Figure 2D), suggesting that the variable mTORC1 activity may be driving the heterogeneous protein synthesis rates across aRVMs. The PE stimulation of protein synthesis required the activation of mTORC1 signaling as non-insulin starved aRVMs maintained high mTORC1 activity and protein translation that completely abolished the PE augmentation of puromycin incorporation (Supplemental Figure 2). Combined, the data show that both mTORC1 activity and protein synthesis are highly heterogeneous in aRVMs and strongly correlated.

**Figure 2.**
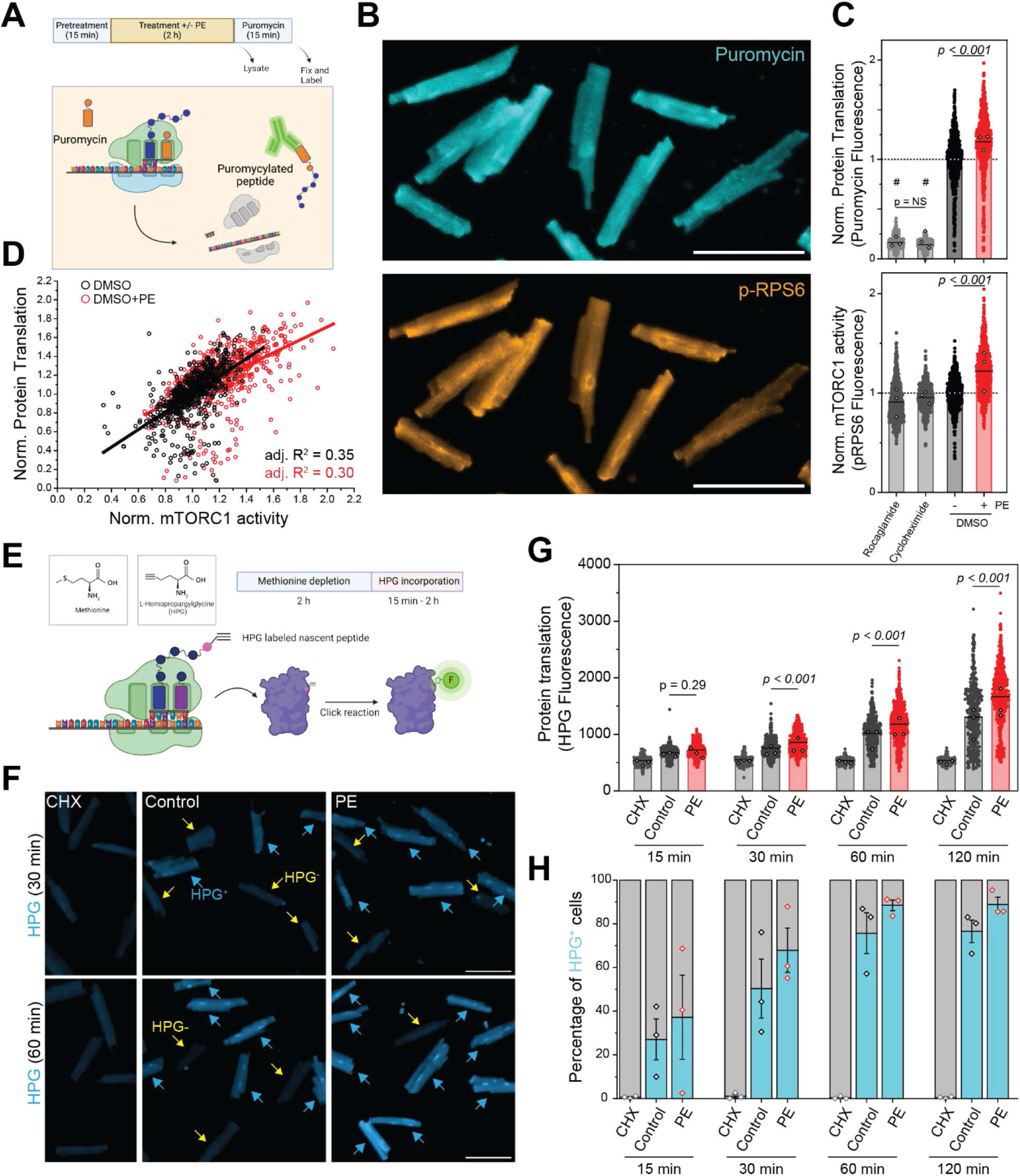
Protein synthesis rate and mTORC1 activity are heterogeneous in isolated aRVMs and stimulated by PE. (A) Experimental protocol for treating aRVMs with inhibitors and PE (10 µM), followed by a 15-minute pulse of puromycin (1 µM). Pretreatment with rocaglamide (3 µM), CHX (18 µM), or any other inhibitor used throughout the study began 15 minutes before stimulation with PE (10 µM, 2 h) and continued through the puromycin treatment. Diagram depicting the mechanism of puromycin incorporation. Puromycin is covalently attached to growing polypeptide chains during the translocation process leading to premature termination and the release of a puromycylated peptide. (B) Representative widefield images of aRVMs labeled with an antibody against puromycin (top) and pRPS6 (bottom). (C) Quantification of relative cellular puromycin fluorescence (top) and pRPS6 fluorescence (bottom). Horizontal line represents the mean of all points. Closed circles represent single-cell measurements and open diamonds represent the mean values from each biological replicate (N = 3 aRVM isolations). Statistical significance was determined by 1-way ANOVA with Bonferroni correction. (D) Correlation between normalized pRPS6 fluorescence and normalized puromycin fluorescence for DMSO and DMSO+PE groups from Figure 2C. Linear fits of all data for each group are shown along with the adjusted R^2^ values for each fit. (E) Diagram of HPG incorporation assay. Inset: Comparison of the structure of methionine and HPG. After a 2 h incubation in methionine free media containing the indicated treatments, aRVMs are incubated in HPG containing methionine free media for the indicated durations. (F) Representative widefield images of HPG labeling in cycloheximide treated (CHX, 18 µM), control, or PE (100 μM) treated aRVMs. aRVMs that incorporate HPG (HPG+) are annotated with cyan arrows, while aRVMs that do not (HPG-) are labeled with yellow arrows. (G) Quantification of the cellular HPG fluorescence (N=3 aRVM isolations). Filled circles represents individual cell measurements and open diamonds represent mean values for each biological replicate. Statistical significance measured using 2-way ANOVA with Bonferroni correction. The measured probabilities of the specific interactions are indicated. (H) Fraction of HPG positive aRVMs calculated from Figure 2G. A threshold was set at the mean + 3 standard deviations of the fluorescence intensities measured from CHX treated aRVMs. The fraction of cells with fluorescence intensity values above the threshold was calculated for each group at each time point. Open diamonds represent mean values for each biological replicate.

Next, we confirmed the heterogeneous protein translation in aRVMs by using an alternative measure of single-cell protein translation with the methionine analog homopropargylglycine (HPG). HPG contains an alkyne moiety that allows for fluorescent labeling using a click chemistry reaction (Figure 2E). Unlike puromycin, HPG does not lead to premature termination and reflects a time integrated measure of protein synthesis over the incubation period. Furthermore, methionine is under-represented across the entire proteome (Supplemental Figure 3) but is obligately incorporated at the start codon; this enrichment at the start codon suggests that incorporation of HPG (a methionine analog) would be biased towards reporting translation initiation.

Considerable heterogeneity in HPG fluorescence was observed. With 30 min incubation of HPG, roughly half of aRVMs displayed robust HPG incorporation suggesting that these cells are translationally active (HPG+). Unexpectedly, the remaining cells displayed low HPG fluorescence intensities comparable to CHX treated aRVMs (Figure 2F), suggesting that these translationally silent cells (HPG-) were not actively producing proteins. Increasing the HPG incubation time to 60 min increased not only the overall fluorescence intensity but also the fraction of HPG+ cells (cyan arrows), suggesting that more translationally silent cells began synthesizing proteins during this time. Quantification of cellular HPG fluorescence confirmed that mean HPG fluorescence increases with increasing incubation durations, which was completely inhibited by CHX treatment (Figure 2G). Surprisingly, treatment of aRVMs with PE increased the mean HPG fluorescence not by uniformly increasing the fluorescence intensity of all cells but by shifting the distribution of cells towards a higher HPG fluorescence (Figure 2G). To quantify this distribution shift, we calculated the fraction of translationally active cardiomyocytes by counting the number of cells displaying HPG fluorescence intensities greater than the mean + 3 standard deviations of CHX treated cells. More cardiomyocytes appeared translationally active with increasing HPG incubation times, and PE augmented the fraction of HPG+ cells (Figure 2H). Combined, these results emphasize that translational activity is highly heterogeneous in aRVMs with individual cells incorporating HPG at different rates.

Hypertrophic stimulation increased the fraction of translationally active cells, supporting an augmentation of protein synthesis potentially through increased translation initiation. Overall, aRVMs recapitulate the heterogeneous mTORC1 activity observed *in vivo*, which is associated with heterogeneous protein synthesis rates *in vitro*, and hypertrophic stimulation with PE recruits more cells into a high mTORC1 activity state to globally augment protein synthesis.

### mTOR inhibition strongly suppresses translation in aRVMs through rapamycin insensitive pathways

While translation initiation is generally thought to be the rate-limiting step in protein synthesis, previous work in adult cardiomyocytes have questioned the importance of this process in driving protein translation ^29^. To establish causal links between mTORC1 activity and protein synthesis, and to dissect the individual contributions of downstream mTORC1 substrates that regulate protein synthesis, we repeated the experiments in the presence of mTOR inhibitors, rapamycin and torin1. Rapamycin is an allosteric inhibitor that forms a complex with FKBP12 to bind mTORC1 and sterically block the phosphorylation of certain substrates ^19,30^. S6K activation upstream of RPS6 phosphorylation is particularly sensitive to inhibition by rapamycin. Torin1 is an ATP-competitive kinase inhibitor that blocks phosphorylation of all downstream mTOR substrates ^31^. As expected, both rapamycin and torin1 abolished pRPS6 labeling (Figure 3A, bottom). mTOR inhibition reduced protein synthesis rates by 10% and 30% with rapamycin and torin1 treatments, respectively (Figure 3A, top). We next examined the effects of hypertrophic stimulation on both mTORC1 activity and protein synthesis rates in aRVMs. We treated aRVMs with PE in the presence of mTOR inhibitors rapamycin and torin1 and both treatments fully blocked the PE induced augmentation of protein synthesis, despite a subtle increase in pRPS6. These results demonstrate that mTORC1 activation is required for PE induced augmentation of protein synthesis.

**Figure 3.**
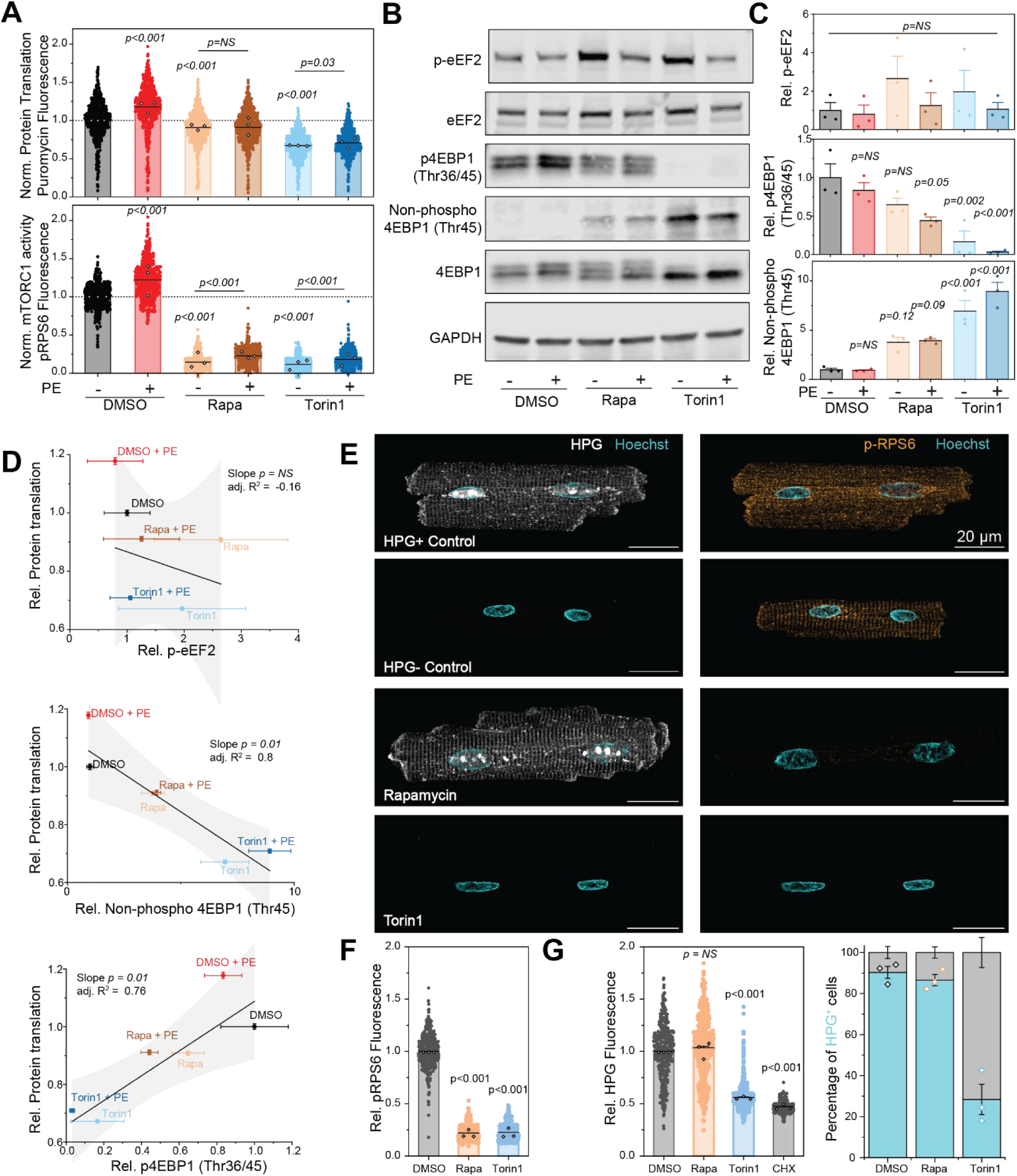
Protein translation in aRVMs is strongly mTOR dependent but largely rapamycin insensitive. (A) Quantification of relative cellular puromycin fluorescence (top) and pRPS6 fluorescence (bottom). Pretreatment with rapamycin (Rapa, 100 nM) or torin1 (250 nM) began 15 minutes before stimulation with PE (10 µM, 2 h) and continued through the puromycin treatment as described in Figure 2A. Horizontal line represents the mean of all points. Closed circles represent single-cell measurements and open diamonds represent the mean values from each biological replicate (N = 3 aRVM isolations). Statistical significance was determined by 1- way ANOVA with Bonferroni correction. Individual comparisons with DMSO are indicated above data and specific comparisons are denoted by horizontal lines. (B) Representative western blots of the indicated mTORC1 substrate measured from lysates prepared with the indicated treatments. Samples were treated with the identical protocol as in A except the puromycin treatment was omitted. (C) Quantification of western blots. All phosphorylation specific bands were normalized to the total substrate. Open diamonds represent the mean values from each biological replicate (N = 3 aRVM isolations). Statistical significance was determined by 1-way ANOVA with Bonferroni correction. (D) Correlations between the bulk abundance of each phosphorylation site in (C) with the mean puromycin incorporation measured in (A). Mean ± standard error. Black lines represent linear fits of the data, and the 95% confidence intervals are highlighted in gray. (E) Representative Airyscan images of aRVMs treated with DMSO (control), rapamycin (100 nM), or Torin1 (250 nM). Immunofluorescence staining for HPG (top) and pRPS6 (bottom) from the same cardiomyocyte are shown with the nuclear dye Hoechst. (F) Quantification of pRPS fluorescence normalized to the DMSO control for each biological replicate. Filled circles represents individual cell measurements and open diamonds represent mean values for each biological replicate. Statistical significance was determined by 1-way ANOVA with Bonferroni correction. (G) Quantification of the HPG fluorescence (1h incubation, left) and the calculated fraction of HPG+ aRVMs (right). Filled circles represents individual cell measurements and open diamonds represent mean values for each biological replicate. Statistical significance was determined by 1-way ANOVA with Bonferroni correction.

We next examined substrate phosphorylation downstream of mTORC1 activation, with particular emphasis on eEF2 phosphorylation downstream of S6K activation, and 4EBP1 phosphorylation, which is directly downstream of mTORC1. Phosphorylation of eEF2 displayed non-significant trends upward with mTORC1-S6K inhibition and modestly downwards with PE stimulation (Figure 3B&C). Importantly, eEF2 phosphorylation was not significantly correlated with global protein synthesis in aRVMs (Figure 3D), suggesting that translation elongation rate does not strongly contribute to global protein synthesis rates in aRVMs. In contrast, 4EBP1 phosphorylation at Thr36/45 was only mildly responsive to rapamycin treatment but was completely abolished with torin1 treatment. This observation was confirmed using an antibody recognizing the non-phosphorylated 4EBP1 (Thr45), which showed a reciprocal response: the non-phosphorylated band was undetectable under baseline conditions but gradually appeared with increasing mTOR inhibition with rapamycin and torin1 treatments. The combination of the two antibodies demonstrates that graded mTOR inhibition resulted in an incremental 4EBP1 dephosphorylation at Thr36/45 that correlated strongly with increasing protein synthesis inhibition (Figure 3D) suggesting that translation initiation is likely driving global protein synthesis rates.

The large fraction of torin1 insensitive puromycin incorporation suggested that the measured protein translation with mTOR inhibition may partly reflect residual ribosomes remaining on long muscle-specific transcripts after the 2-hour torin1 pretreatment. Puromycin incorporation is biased towards reporting translation elongation given its mechanism of action (Figure 2A), which may downplay the contribution of mTORC1 activity to protein translation. To further evaluate the contribution of mTORC1 signaling to protein synthesis, we measured HPG labeling with mTOR inhibition with rapamycin or torin1. We again observed a mixed population of HPG+ and HPG- cells under DMSO control conditions (Figure 3E). Both rapamycin and torin1 abolished pRPS6 labeling as before (Figure 3E bottom row, 3F) and similar results were observed when cardiomyocytes were stained with an antibody against p- mTOR (Supplemental Figure 4A&B). However, mTOR inhibition resulted in divergent HPG labeling patterns with rapamycin treatment having no effect on HPG labeling while torin1 strongly suppressed HPG labeling (Figure 3E&G). Quantification of HPG fluorescence revealed that rapamycin treated cells did not alter HPG fluorescence and displayed a similar fraction of translationally active HPG+ cells compared to DMSO treated control cells (Figure 3E-G). In contrast, mTOR inhibition with torin1 strongly shifted the distribution of HPG fluorescence intensities down to levels observed in CHX treated cells, resulting in a significantly smaller fraction of translationally active HPG+ cells (Figure 3E-G and Supplemental Figure 4C).

Treatment with LY294002 to inhibit PI3K activation upstream of mTORC1 activation, also led to a strong suppression of HPG labeling (Supplemental Figure 4C). Rocaglamide completely abolished HPG incorporation, indicating that HPG labeling is strongly dependent on cap-dependent translation initiation (Supplemental Figure 4C). Combined, these results demonstrate that protein synthesis is largely driven by rapamycin insensitive substrates downstream of mTOR and implicates translation initiation as the primary driver of global protein synthesis in aRVMs.

### Differential phosphorylation of 4EBP1 at Thr36/45/69 drive heterogeneous protein synthesis in aRVMs

Given the strong dependence of protein synthesis on 4EBP1 phosphorylation, we hypothesized that the heterogeneous mTORC1 activity would lead to differential phosphorylation of 4EBP1 across aRVMs, resulting in the variable translation rates. To test this, we co-labeled cardiomyocytes with puromycin and phospho-antibodies targeting the canonical mTORC1 dependent phosphorylation sites Thr36/45 and Thr69. We also determined the PE, rapamycin, and torin1 dependence of these phosphorylation sites. Labeling aRVMs with the p4EBP1 (Thr36/45) antibody recapitulated the cell-to-cell heterogeneity observed with pRPS6 (Figure 4A, top) and provided additional evidence of heterogeneous mTORC1 activity in aRVMs. The heterogeneity in 4EBP1 phosphorylation at Thr36/45 was confirmed by labeling aRVMs with the non-phosphorylated 4EBP1 (Thr45) antibody (Figure 4A, bottom). While changes in phosphorylation with PE at Thr36/45 could not be detected in bulk lysates (Figure 3B), single myocyte imaging revealed an increase in 4EBP1 phosphorylation at Thr36/45 and a corresponding decrease in the non-phospho 4EBP1 (Thr45) upon PE stimulation (Figure 4B). Rapamycin had no effect on phosphorylation at Thr36/45, consistent with the reported rapamycin resistance of these priming sites ^32^. mTOR inhibition with torin1 strongly suppressed phospho 4EBP1 (Thr36/45) and strongly increased non-phospho 4EBP1 (Thr45) labeling, confirming the mTOR dependence of phosphorylation at these sites. Overall, the magnitude of phosphorylation at Thr36/45 was strongly correlated with puromycin labeling (Figure 4C), further demonstrating that underlying heterogeneity in mTORC1 activity drive heterogeneous 4EBP1 phosphorylation and subsequent translation initiation in aRVMs.

**Figure 4.**
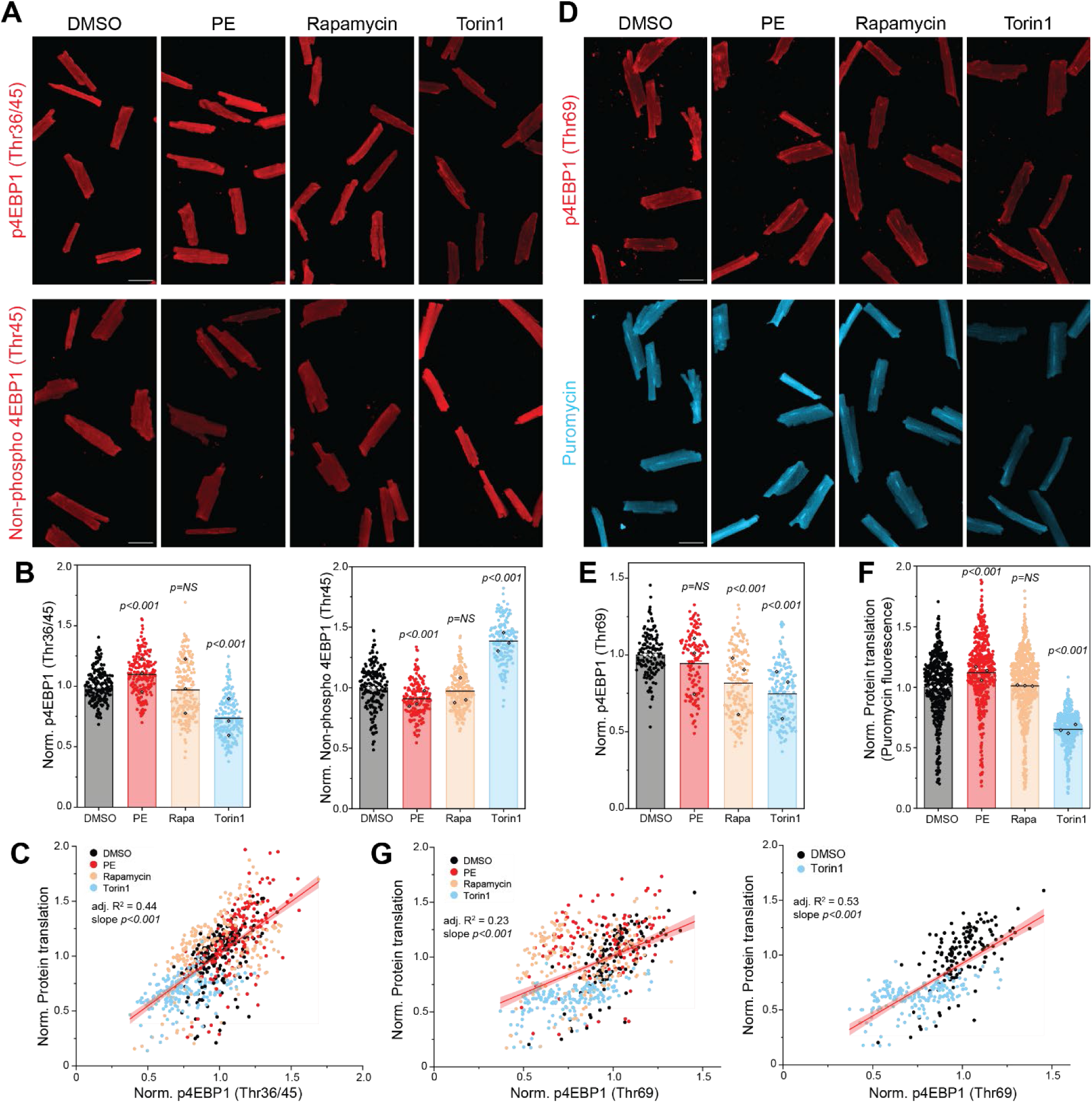
Differential phosphorylation of 4EBP1 at Thr36/45/69 drives heterogeneous protein synthesis in aRVMs. (A) Representative tile scan images of aRVMs co-labeled with puromycin and p4EBP1 (Thr36/45) on top or non-phospho 4EBP1 (Thr45) on bottom and treated with the indicated pharmacological agent. Scale bar represents 50 µm. (B) Quantification of single cell fluorescence intensities of the indicated phospho-specific antibody. Close circles represent individual cells, and open diamonds represent the mean values from each biological replicate. N = 3 aRVM isolations. (C) Correlation between p4EBP1 Thr36/45 and puromycin fluorescence. Closed circles represent individual cell measurements. A linear fit of the total data are shown in red. The shaded pink regions represent the 95% confidence intervals of the fit. (D) Representative tile scan images of aRVMs co-labeled with puromycin and p4EBP1 (Thr69) on top and the same cells displaying the puromycin intensity on bottom and treated with the indicated pharmacological agent. Scale bar represents 50 µm. (E) Quantification of single cell fluorescence intensity of p4EBP1 (Thr69) and (F) combined puromycin fluorescence intensities co-labeled with each phospho-specific antibody. Close circles represent individual cells, and open diamonds represent the mean values from each biological replicate. N = 3 aRVM isolations. (G) Correlation between p4EBP1 Thr69 and puromycin fluorescence for all 4 groups (left) and DMSO and Torin1 only (right).

We next examined the PE and mTOR dependence of 4EBP1 phosphorylation at Thr69 (Figure 4D). Labeling for p4EBP1 Thr69 was suppressed with both rapamycin and torin1 treatment (Figure 4E), consistent with the reported mTOR dependence and rapamycin sensitivity of this site ^33^. Unexpectedly, PE stimulation did not alter phosphorylation at Thr69, despite increasing protein synthesis (Figure 4F), demonstrating that an alternative mechanism drives the augmented translation. p4EBP1 Thr69 was also correlated with puromycin incorporation (Figure 4G) but the relationship between rapamycin sensitive mTORC1 activity and protein synthesis was confounded by the deviations in signaling following rapamycin and PE treatments.

Examination of this relationship under basal conditions and following torin1 treatment revealed a stronger positive correlation suggesting that hyperphosphorylation of 4EBP1 at Thr69 by canonical mTORC1 signaling also contributes to the basal heterogeneity of protein translation observed in aRVMs, but may not contribute to augmented protein synthesis upon hypertrophic stimulation. This antibody did not detect bands in western blots with aRVM lysates. The combined results demonstrate that heterogeneous mTORC1 dependent 4EBP1 phosphorylation at Thr36/45 and Thr69 are the primary drivers of heterogeneous protein synthesis rates across cardiomyocytes at baseline.

### mTORC1 independent, MEK-ERK dependent phosphorylation of 4EBP1 Ser64 partly mediates protein synthesis augmentation following PE stimulation

Given our observation that hypertrophic stimulation with PE is not mediated through canonical mTORC1 dependent 4EBP1 Thr69 phosphorylation, we next examined 4EBP1 phosphorylation at Ser64, which is robustly detected following pressure-overload induced cardiac hypertrophy ^34,35^. PE treatment of aRVMs strongly induced phosphorylation at this site in western blots (Figure 5A) suggesting that this site may act as a putative mechanism to augment protein translation. Unfortunately, this antibody was not amenable to detection by immunofluorescence due to high non-specific labeling (Supplemental Figure 5). Although 4EBP1 Ser64 is known to be strongly rapamycin sensitive in non-myocytes ^32^, we unexpectedly observed that Ser64 phosphorylation is unaffected by rapamycin or torin1 pretreatment (Figure 5A), suggesting that this site may not be regulated by canonical mTORC1 signaling in aRVMs. We next tested whether Ser64 phosphorylation in aRVMs followed the hierarchical phosphorylation model that required prior phosphorylation at the priming sites Thr36/45. To do so, aRVMs were treated with Torin1 for 2 hours to abolish Thr36/45 phosphorylation and then stimulated with PE for 1 hour (Figure 5B). Under these conditions, PE was still able to phosphorylate 4EBP1 Ser64 in the absence of detectable Thr36/45 phosphorylation, indicating that prior phosphorylation at Thr36/45 is not necessary for phosphorylation at Ser64 in aRVMs, demonstrating a substantial deviation from the canonical mTORC1 dependent hierarchical phosphorylation model of 4EBP1. These results suggested that an alternative kinase phosphorylates 4EBP1 at this site independently of mTORC1 in cardiomyocytes.

**Figure 5.**
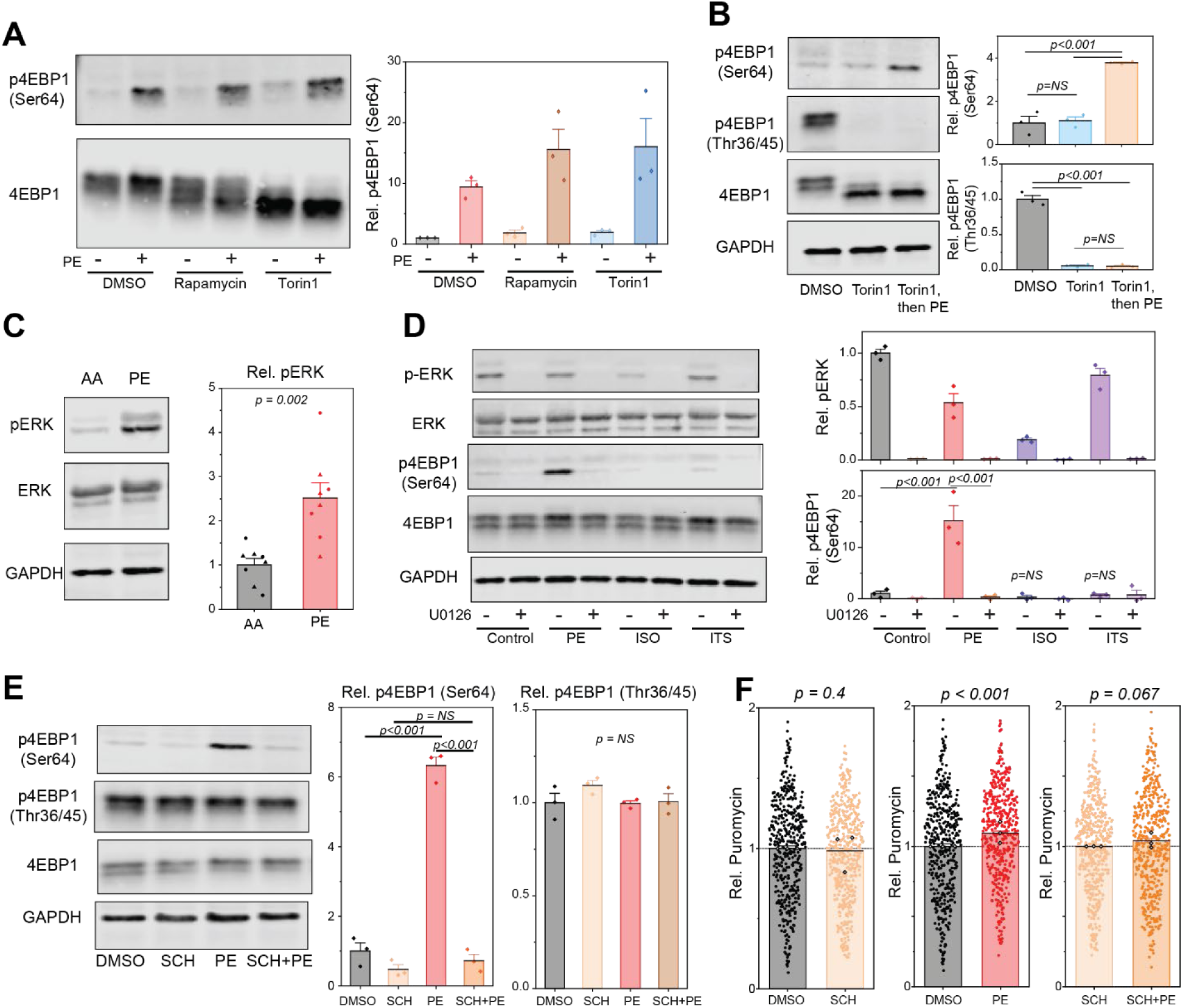
ERK phosphorylates 4EBP1 at Ser64 in aRVMs. (A) mTOR inhibition does not suppress PE induced phosphorylation of 4EBP1 at Ser64. Western blots of lysates prepared from aRVMs treated with mTOR inhibitors, rapamycin (100 nM) or torin1 (250 nM) and PE (10 µM). The treatment protocol is described in Figure 2A. The quantification in A, B, D and E represent the mean + standard error of phosphorylated band intensities normalized to their respective total substrates. Closed diamonds represent mean values from each biological replicate N=3 aRVM isolations. (B) Western blots of lysates prepared from aRVMs treated with Torin1 (250 nM) for 2h or Torin1 for 2h followed by 1h PE treatment. (C) Western blots of lysates prepared from 4h AA or PE hearts blotted for ERK activation. Closed circles represent individual male animals and closed triangles represent individual female animals. Mean + Standard Error. (D) Western blots of lysates prepared from aRVMs treated with the MEK inhibitor U0126 (10 µM) and either PE (10 µM), isoproterenol (ISO, 100 nM), or ITS (1X) following the treatment protocol in Figure 2A. (E) Western blots of lysates prepared from aRVMs treated with the ERK inhibitor SCH772984 (250 nM) and PE (10 µM). (F) Quantification of puromycin incorporation in aRVMs pre-treated with SCH772984 for 15 minutes, followed by PE (10 µM) for 2 hours. (right) The puromycin fluorescence was normalized to either the DMSO or SCH control. Closed circles represent individual cells, and open diamonds represent the mean value of each biological replicate (N=3 aRVM isolations).

Alternative kinases such as CDK1 are known to phosphorylate 4EBP1 at Ser64 during mitosis ^36^, when 4EBP1 is hypophosphorylated ^37^. However, it is unlikely that these mechanisms persist in post-mitotic cardiomyocytes. While 4EBP1 phosphorylation by ERK has not been demonstrated in cells, ERK has been used to phosphorylate multiple 4EBP1 phosphorylation sites including Ser64 in purified proteins ^38^. We hypothesized that ERK may be the putative kinase that regulates 4EBP1 at Ser64. To test this, we first confirmed that PE stimulation *in vivo* leads to ERK activation. A robust phosphorylation of p42/44 ERK at Tyr 202/204 was observed in our 4h PE stimulation model (Figure 5C), confirming ERK activation with hypertrophic stimulation. Next, we tested the specificity of p4EBP1 Ser64 to PE stimulation and the MEK-ERK dependence of this phosphorylation site. Treatment of cardiomyocytes with PE led to Ser64 phosphorylation, which was not observed with the β-adrenergic receptor agonist isoproterenol or insulin, suggesting a distinct signaling pathway activated with PE (and pressure-overload ^34,35^) (Figure 5D). Pre-treatment of aRVMs with the non-selective MEK inhibitor U0126 completely abolished ERK phosphorylation and PE induced 4EBP1 phosphorylation at Ser64. Similarly, the selective ATP-competitive ERK1/2 inhibitor, SCH772984 (SCH) ^39^ also inhibited phosphorylation of 4EBP1 at Ser64 in response to PE treatment (Figure 5E). Importantly, SCH treatment had no effect on 4EBP1 phosphorylation at Thr36/45, confirming that the acute treatment did not alter mTORC1 function. Combined, the data demonstrate that PE induced 4EBP1 phosphorylation at Ser64 is MEK-ERK dependent in aRVMs.

Next, the functional role of 4EBP1 phosphorylation at Ser64 was examined by measuring puromycin incorporation in SCH treated cardiomyocytes. While ERK inhibition with SCH had no effect on basal protein synthesis rates (Figure 5F), pre-treatment with SCH abolished the PE induced increase in protein synthesis rates, demonstrating a direct role of ERK in promoting translation initiation and protein synthesis in cardiomyocytes during hypertrophic stimulation.

### PE stimulated hypertrophy *in vivo* is mediated through mTORC1 dependent phosphorylation of 4EBP1 at Thr36/45 and ERK dependent phosphorylation at Ser64

Having separated the mTOR and ERK dependent translational control mechanisms and their contributions to protein synthesis rate in aRVMs, we went back to our pre-hypertrophy model to determine which of these effectors are active under basal conditions and after PE stimulation *in vivo*. Similar responses were observed in both male and female animals treated with PE. Western blotting of ventricular lysates revealed a robust increase in the pRPS6 (Figure 6A), consistent with the activation of the mTORC1 signaling pathway observed in tissue sections (Figure 1). Total levels of ribosomal protein S6 (RPS6) were mildly upregulated, suggesting that ribosome biogenesis was initiated within 4 hours and a modest increase in ribosome abundance at this time point. Given the early activation of ribosome biogenesis, we tested if inhibition of transcription to block ribosome biogenesis could suppress PE induced augmentation of protein translation in aRVMs. Actinomycin D blocks transcription by inhibiting RNA polymerases, including RNA Pol I and III that are essential for rDNA transcription and ribosome biogenesis ^40^. We observed that actinomycin D had no effect on basal puromycin incorporation but blunted the PE induced augmentation of protein synthesis (Supplemental Figure 6), suggesting that some ribosome biogenesis may moderately contribute to the acute (<4 h) PE induced cardiac hypertrophy.

**Figure 6.**
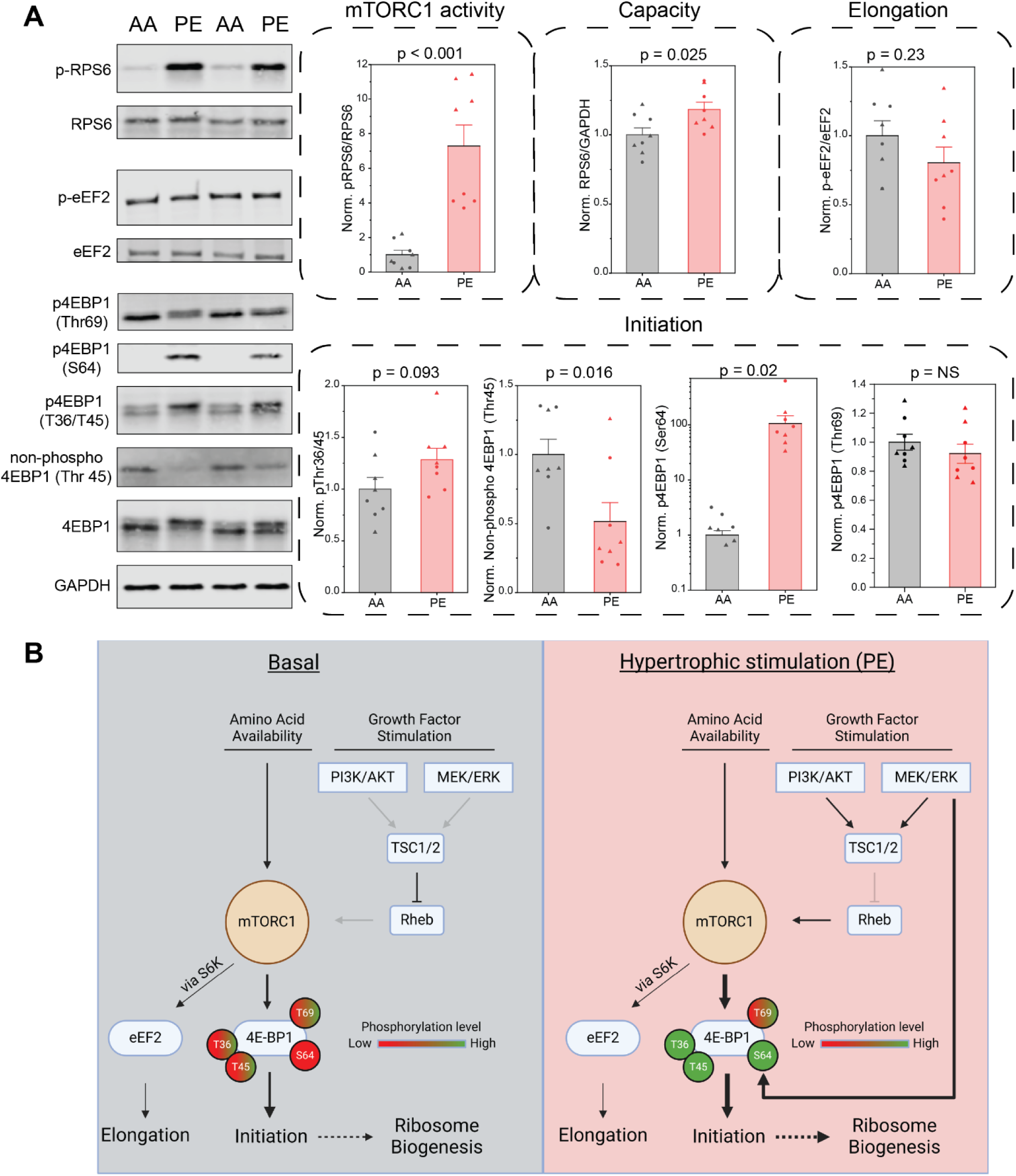
Translational control contributions by mTORC1 and ERK *in vivo*. (A) Western blots for mTORC1 activation, eEF2 phosphorylation, and 4EBP1 phosphorylation. Quantification of the western blots and the corresponding translational control mechanisms are grouped together. Closed circles represent individual male animals and closed triangles represent individual female animals. Mean + Standard Error. N = 8 (4 male, 4 female) AA, 8 (4 male, 4 female) PE. (B) Schematic depicting the relative contributions of translational control mechanisms to protein synthesis under basal conditions and after hypertrophic stimulation. The thickness of the arrows indicates the relative contribution of the specific signaling pathway. Graded levels of phosphorylation of the indicated 4EBP1 sites are color coded.

eEF2 phosphorylation was slightly but non-significantly reduced in PE treated animals, suggesting that regulation of translation elongation plays a minor role in hypertrophic augmentation of translation. In contrast, we observed dramatic changes in the phosphorylation of the translation initiation repressor 4EBP1. We observed a clear band for non-phosphorylated 4EBP1 (Thr45) under basal conditions that was lost following PE stimulation and a reciprocal trend towards increased p4EBP1 (Thr36/45) following PE stimulation (Figure 6A). Consistent with our observations in aRVMs, phosphorylation of 4EBP1 at Thr69 was unaltered with PE despite the strong mTORC1 activation and, PE stimulation strongly induced phosphorylation of 4EBP1 at Ser64. Notably, the abundance of p4EBP1 (Ser64) was strongly correlated with the magnitude of ERK activation, but not mTORC1 activation, in support of the ERK dependence of this phosphorylation site (Supplemental Figure 7).

Overall, the *in vivo* data are consistent with the concerted activation of protein synthesis augmentation through graded increases in translation initiation via 4EBP1 phosphorylation mediated by two distinct yet converging pathways (Figure 6B). Combined, our data indicate that the primary drivers of protein synthesis upon PE stimulation are 1) mTORC1 dependent 4EBP1 phosphorylation at Thr36/45 and 2) ERK dependent 4EBP1 phosphorylation at Ser64. Consequently, the increased translation initiation leads to a rapid increase in ribosome abundance that is detectable within hours after hypertrophic stimulation.

## DISCUSSION

### mTORC1 heterogeneity in the heart

In this study, we performed a single-cell investigation of translational control mechanisms in adult cardiomyocytes. Our results reveal a surprisingly high heterogeneity in mTORC1 activity that results in heterogeneous protein synthesis rates across cardiomyocytes, and which is driven primarily by differential 4EBP1 phosphorylation. Additionally, our findings demonstrate that hypertrophic stimulation drives augmented protein synthesis by recruiting more cardiomyocytes into a high mTORC1 activity state to phosphorylate 4EBP1 at Thr36/45, and by promoting MEK-ERK dependent phosphorylation of 4EBP1 at Ser64.

We find that the heart has a low basal level of mTORC1 activity, indicated by the predominance of low pRPS6 labeled cardiomyocytes (Figure 1C&D) and robust detection of non-phosphorylated 4EBP1 (Thr 45) at baseline (Figure 6A). External factors such as nutrition status and time of day can influence basal mTORC1 activity *in vivo*. Cardiac mTORC1 activity shows circadian fluctuations peaking during the light phase when rodents are at rest ^41^. Since all of our cardiac tissues were collected during the daytime and animals were not fasted, we estimate that these basal levels reflect the upper range of mTORC1 activity in the heart under non-stressed conditions. Thus, the low mTORC1 activity is a driver for the low basal translation rate in the heart ^3^, in addition to other reported factors including the relatively low expression of translational machinery ^42^ and PABPC1 ^43^.

By operating at the low end of mTORC1 activity, the heart maintains a large reserve capacity to adapt acutely to external stimuli. This regulation was clearly evident with a bolus PE stimulation, which robustly augmented global mTORC1 activity *in vivo* by recruiting more cardiomyocytes into a high mTORC1 activity state. Yet, the same regulatory mechanisms may be constitutively operating at a local level under basal conditions. Seminal work by Cooper et. al. demonstrated that cardiac hypertrophy is regionally regulated by load: in the same heart, an unloaded papillary muscle can undergo atrophy while the remaining pressure overloaded heart grows ^44^. Extrapolating this concept further to the single cardiomyocyte level, local differences in load, amino acid availability, metabolism, or growth factor signaling may stimulate mTORC1 activation in individual cardiomyocytes to create the observed mosaic pRPS6 staining. Such a model may allow cardiomyocytes to acutely respond to their local environment, resulting in a variety of cell states that are difficult to capture with bulk assays. While this study primarily focused on the downstream anabolic pathways, the observed heterogeneous mTORC1 activity would also be expected to differentially regulate catabolic pathways including autophagy and lysosomal biogenesis in cardiomyocytes given the hierarchical phosphorylation of TFEB ^45^. These observations demonstrate a need to further explore the variability of phenotypes across cardiomyocytes and to carefully consider the acute environmental conditions and stimuli that can shift the cellular distribution of observable phenotypes.

### Regulation of 4EBP1 in the heart

Our results also demonstrate a novel model of translation initiation control through 4EBP1 phosphorylation. We find that the PE specific activation of the MEK-ERK pathway can directly phosphorylate Ser64 on 4EBP1 (Figure 5D) to bypass mTORC1 dependent phosphorylation of Thr69, giving rise to a dual input model of mTORC1 and ERK dependent phosphorylation of 4EBP1 in cardiomyocytes. While PE can induce phosphorylation of Ser64 even when mTORC1 is pharmacologically inhibited (Figure 5A), mTORC1 activation and 4EBP1 Thr36/45 phosphorylation is required for the ERK dependent Ser64 phosphorylation to have a positive effect on translation following PE stimulation (Figure 2A-C), consistent with biochemical studies demonstrating that phosphorylation at Ser64 alone is insufficient to disrupt 4EBP-eIF4E interaction ^20^. Both insulin and PE activate protein synthesis through 4EBP1 phosphorylation, but our results demonstrate that these agonists activate different upstream pathways to augment translation initiation. Insulin drives mTORC1 activation through the canonical pathway (Supplemental Figure 1) leading to 4EBP1 phosphorylation at Thr36/45/69 ^46^ and did not strongly induce phosphorylation of 4EBP1 at Ser64 (Figure 5D). Conversely, phosphorylation of Thr69 is unaltered with PE stimulation despite an increase in mTORC1 dependent Thr36/45 phosphorylation. These observations support the notion that the distal phosphorylation sites on 4EBP1 are functionally redundant for 4EBP1 inactivation and hypertrophic PE stimulation switches the hyperphosphorylated species of 4EBP1 from the predominant mTORC1-dependent pThr36/45/69 proteoform to the mTORC1/ERK-dependent pThr36/45/Ser64 proteoform.

These findings raise several questions. First, although multiple extracellular growth factors including insulin are known to activate ERK, why is PE the only tested agonist that led to significant phosphorylation of 4EBP1 at Ser64 (Figure 5D)? A potential explanation may be that a Gq coupled receptor specific pathway may be the upstream driver of 4EBP1 phosphorylation at Ser64. Following Gq receptor activation by PE or pressure-overload, ERK binds the released Gβγ subunit and undergoes autophosphorylation at Thr188, which drives dimerization and nuclear translocation of ERK to mediate hypertrophic growth ^47^. This pathway is distinct from canonical ERK activation and confers phosphorylation and activation of a subset of ERK functions during cardiac hypertrophy. Additional studies are necessary to determine if 4EBP1 phosphorylation at Ser64 is dependent on the pERK Thr188 dependent pathway.

Second, if maximal protein translation rates can be achieved through mTORC1 activation alone, what function would a secondary ERK pathway play in hypertrophic growth? While transgenic animals lacking ERK activation can still grow in response to pressure overload or pharmacological stressors ^48^, cardiac hypertrophy in the absence of ERK signaling results in eccentric remodeling ^49^, demonstrating that ERK mediates some as yet unknown pathway to control concentric growth. We recently demonstrated that cardiomyocytes rely strongly on microtubule dependent mRNA and ribosome transport to sites of local translation, primarily at the z-disk, intercalated disk, and the perinucleus (Figure 3E and Supplemental Figure 4A), for productive cardiac growth ^21^. While active mTORC1 localizes to lysosomal membranes ^50^, active ERK translocates to the nucleus following PE stimulation ^47^, demonstrating that these kinases may occupy distinct spatial domains within the cardiomyocyte. Thus, nuclear ERK could selectively phosphorylate 4EBP1 bound to nascent transcripts and prime these transcripts for translation upon encountering active mTORC1 in the cytosol. This Ser64 priming mechanism could preferentially reallocate ribosomes away from pre-existing transcripts to nascent transcripts as they are exported out from nuclei. While speculative, future work could explore the ERK-dependent translatome to potentially identify transcripts that are acutely translated to determine the mechanisms that underlie directed growth.

### Study Limitations

This study used specific kinase inhibitors at nanomolar concentrations and short (<2.5 h) treatment times to minimize off-target effects and prevent cellular adaptation to chronic ablation of specific signaling pathways. Although many of these molecules are selective for a few kinases out of the hundreds of known kinases, care should be taken in the interpretation of these results.

Additionally, differences in amino acid concentrations *in vivo* and in the M199 based culture media likely lead to differences in baseline mTORC1 activation. Three key amino acids leucine, arginine, and methionine are known to activate mTORC1 and are present at ∼ 3-fold higher concentrations in M199 media than in human plasma ^51^. Furthermore, AMPK activity, a known inhibitor of mTORC1 kinase ^4^, is likely higher *in vivo* in an actively beating heart than in quiescent isolated cardiomyocytes. Despite these different contexts, and with mTORC1 activity expected to be significantly lower *in vivo* than in aRVMs, we observed consistent regulation of translational control mechanisms following PE stimulation both *in vivo* and *ex vivo*.

## ACKNOWLEDGEMENTS

The authors would like to thank Li Li, Waixing Tang from the Penn Center for Musculoskeletal Disorders Histology Core for their advice in preparing tissue sections for immunohistochemistry, the CDB Microscopy Core (RRID SCR_022373), Katey Stone and Carmen Suay Corredera for assistance in preparing cardiac lysates, and Rani Randell and Taryn Wilson for assistance in isolating cardiomyocytes.

## Non-standard Abbreviations and Acronyms

4EBP1: eukaryotic translation initiation factor 4E binding protein 1
aRVMs: adult rat ventricular myocytes
CHX: Cycloheximide
eEF2: eukaryotic elongation factor 2
eEF2K: eukaryotic elongation factor 2 kinase
ERK: extracellular signal-regulated kinase
HPG: Homopropargylglycine
mTORC1: mammalian target of rapamycin complex 1
PE: phenylephrine
RPS6: Ribosomal protein S6
RocA: Rocaglamide A
S6K: ribosomal protein S6 kinase

---

**Supplemental Figure 1.**
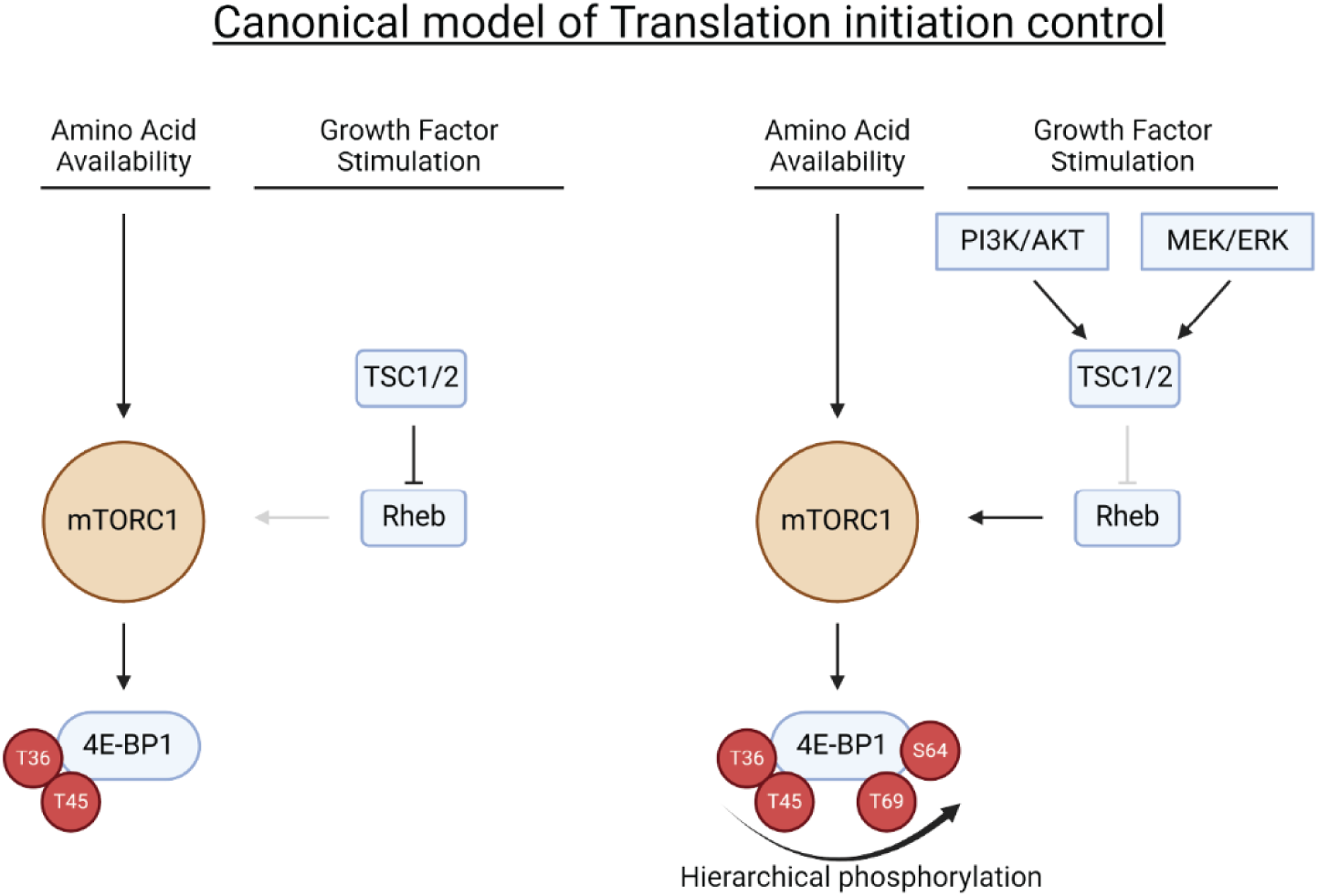
Canonical model of translation initiation regulation. (Left) mTORC1 is active at a basal level when amino acids are sufficiently available but growth factor stimulation is absent. Under this condition, the TSC1/2 complex inhibits Rheb from further activating mTORC1. mTORC1 is thought to preferentially phosphorylate some substrates under these conditions including 4EBP1 at Thr36/45. (Right) With growth factor stimulation (Ex. insulin/IGF-1 stimulation of the PI3K/AKT pathway or adrenergic receptor activation of the MEK/ERK pathway), TSC1/2 becomes phosphorylated and inactivated, thereby relieving the inhibition of Rheb. Rheb can then further activate mTORC1 signaling, which is thought to promote further phosphorylation of mTORC1 substrates including 4EBP1 at Thr69 followed by Ser64.

**Supplemental Figure 2.**
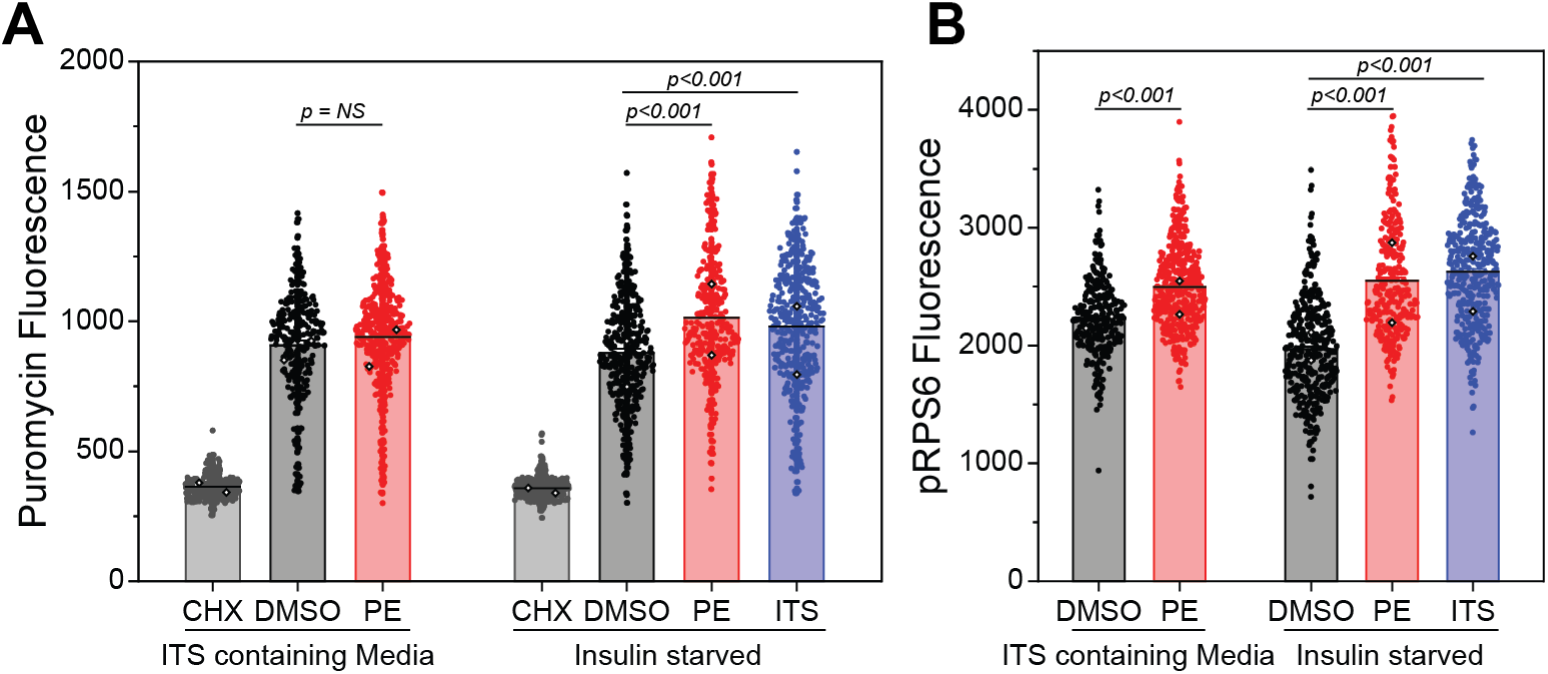
PE cannot stimulate protein synthesis in the presence of saturating insulin concentrations. aRVMs were cultured overnight in media with or without insulin-transferrin-selenium (ITS). The cells were stimulated with 10 µM PE or 1X ITS for 2 hours and (A) puromycin incorporation and (B) RPS6 phosphorylation were measured. Filled circles represents individual cell measurements and open diamonds represent mean values for each biological replicate (N = 2 aRVM isolations). Statistical significance was determined by 1-way ANOVA with Bonferroni correction.

**Supplemental Figure 3:**
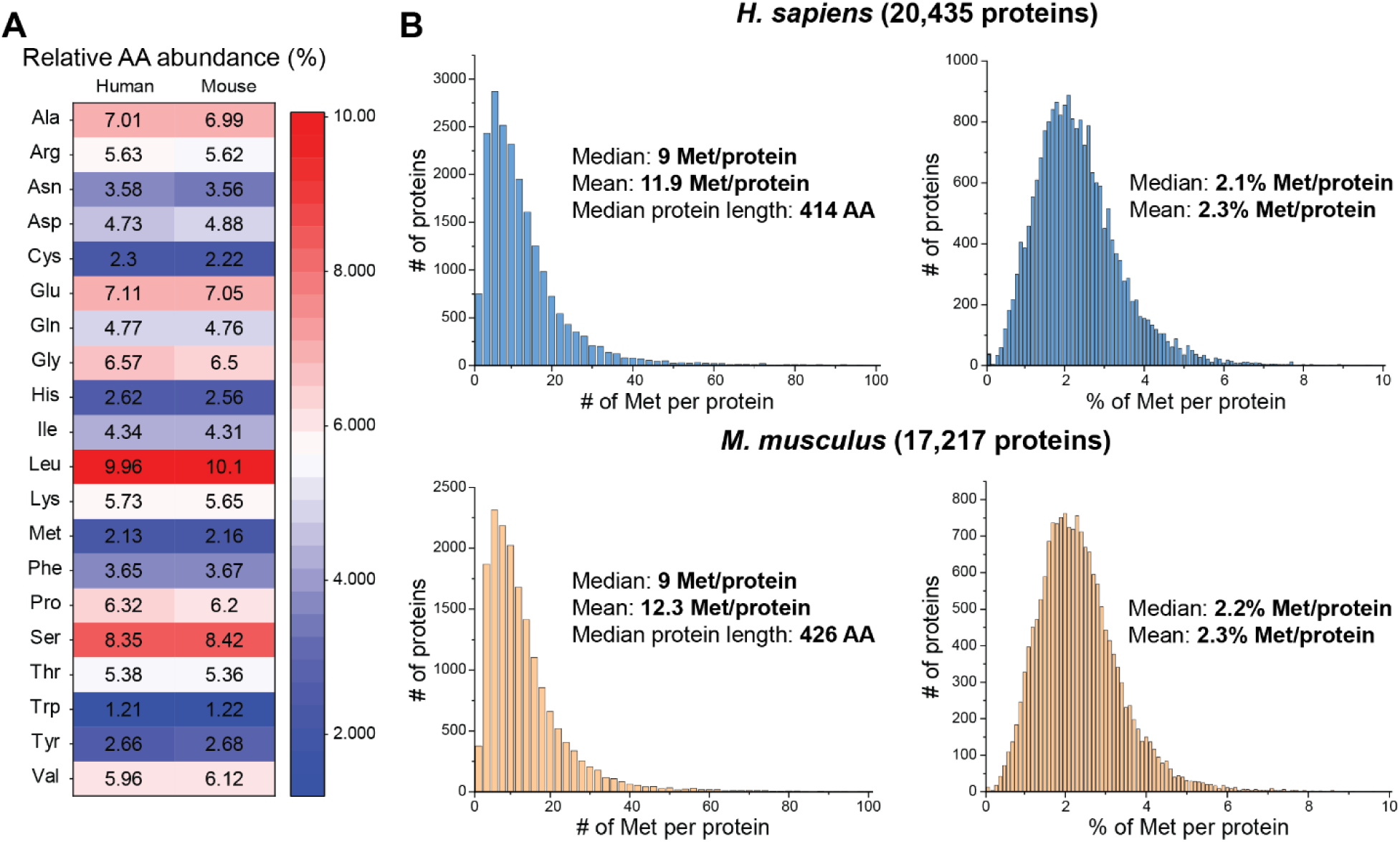
Amino acid usage across human and mouse proteomes. (A) Heatmap of mean abundances of each amino acid in human and mouse proteomes. (B) Histograms representing the number of methionines per protein in human (top) and mouse (bottom) proteomes (left). Histograms representing the relative abundance of methionines per protein in human and mouse proteomes (right).

**Supplemental Figure 4.**
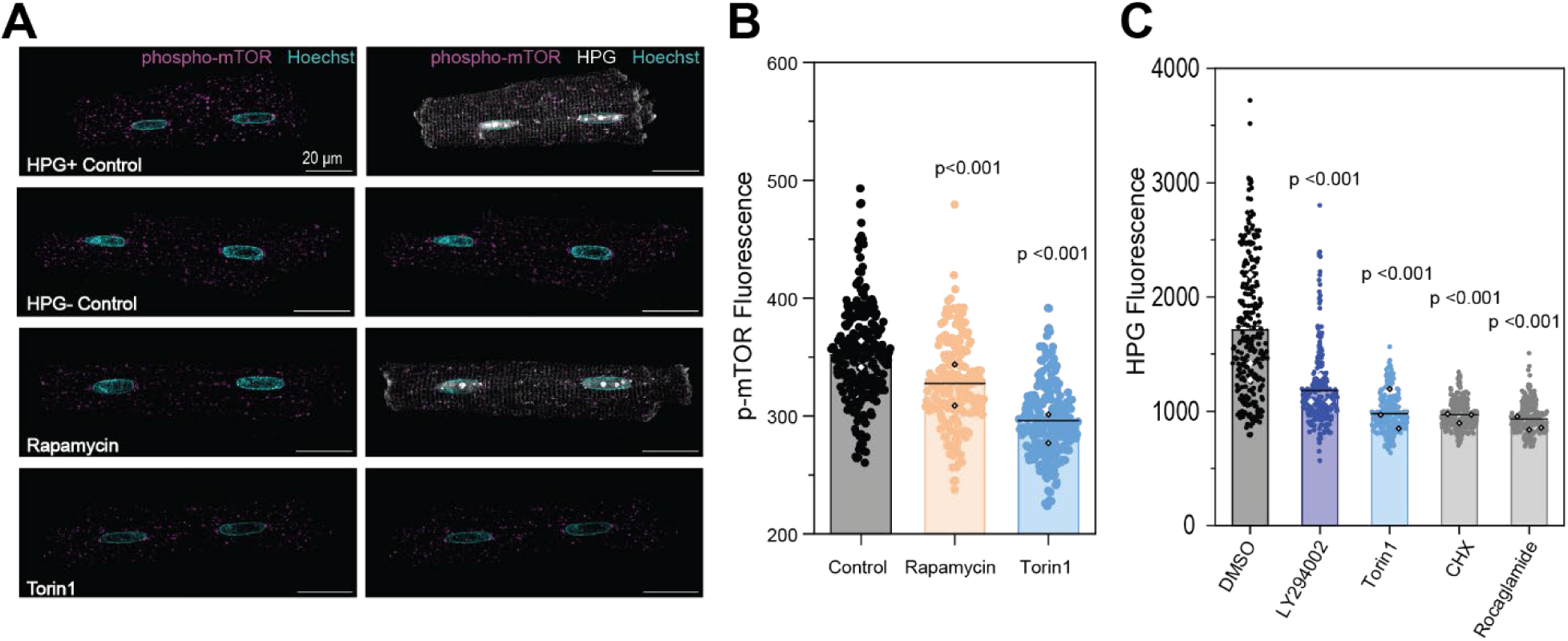
mTOR regulation of HPG labeling. (A) Representative Airyscan confocal images of cardiomyocytes labeled with HPG and phospho-mTOR. (B) Quantification of phospho-mTOR fluorescence from the indicated treatment. Each dot represents a single cell, measured from widefield images. (N=2 aRVM isolations) (C) Quantification of HPG fluorescence with the indicated treatments. Filled circles represents individual cell measurements and open diamonds represent mean values for each biological replicate (N = 3 aRVM isolations). Statistical significance was determined by 1-way ANOVA with Bonferroni correction.

**Supplemental Figure 5.**
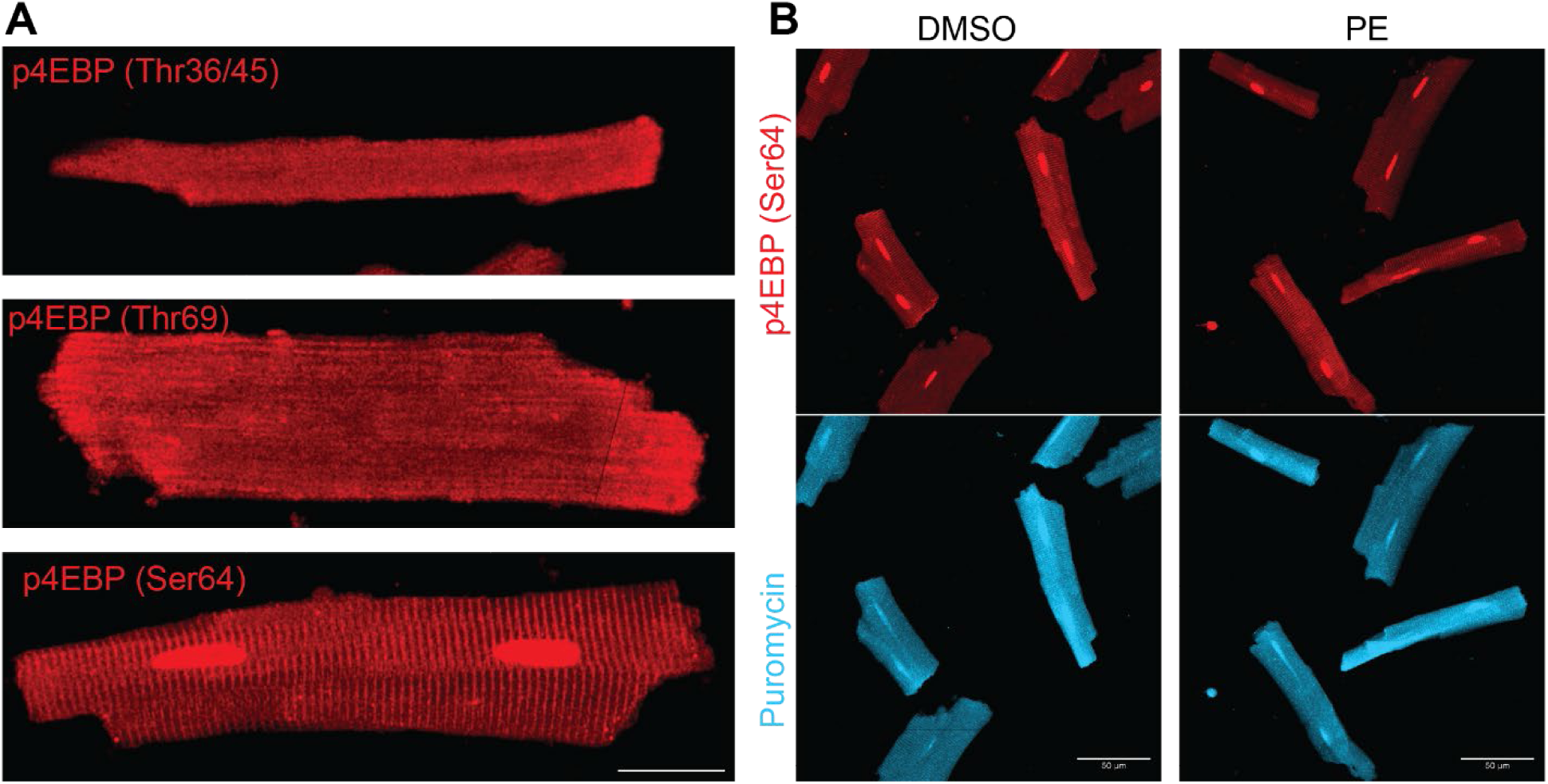
p4EBP1 Ser64 antibody displays non-specific labeling. (A) Zoomed representative images of aRVMs stained for the indicated p4EBP1 antibody. p4EBP1 (Ser64) antibody labeling displays strong intranuclear and z-disk labeling that is distinct from the cytosolic labeling observed with the other 4EBP1 antibodies. Scale bar represents 20 µm. (B) p4EBP1 (Ser64) displays no response to PE (10 µM, 2h) stimulation, demonstrating that this antibody is not suitable for quantitative immunofluorescence.

**Supplemental Figure 6.**
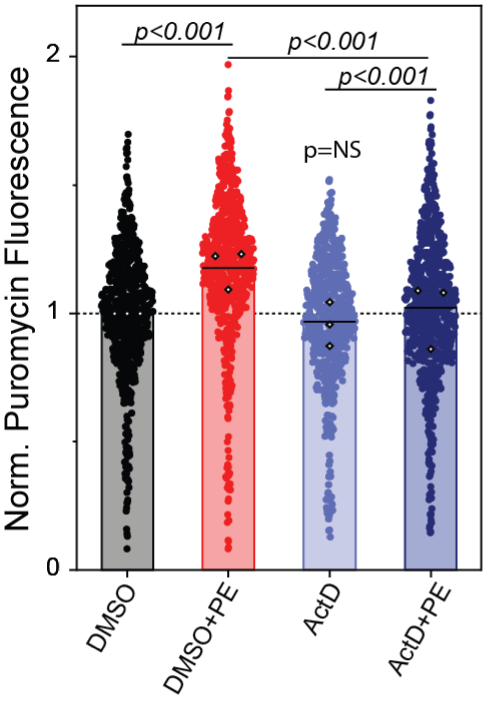
Inhibition of transcription with Actinomycin D (ActD) partially inhibits PE induced protein synthesis augmentation. aRVMs were pre-treated with DMSO or actinomycin D (ActD, 2.5 µM) for 15 minutes before PE treatment (10 µM, 2h). Filled circles represents individual cell measurements and open diamonds represent mean values for each biological replicate (N = 3 aRVM isolations). Statistical significance was determined by 1-way ANOVA with Bonferroni correction.

**Supplemental Figure 7.**
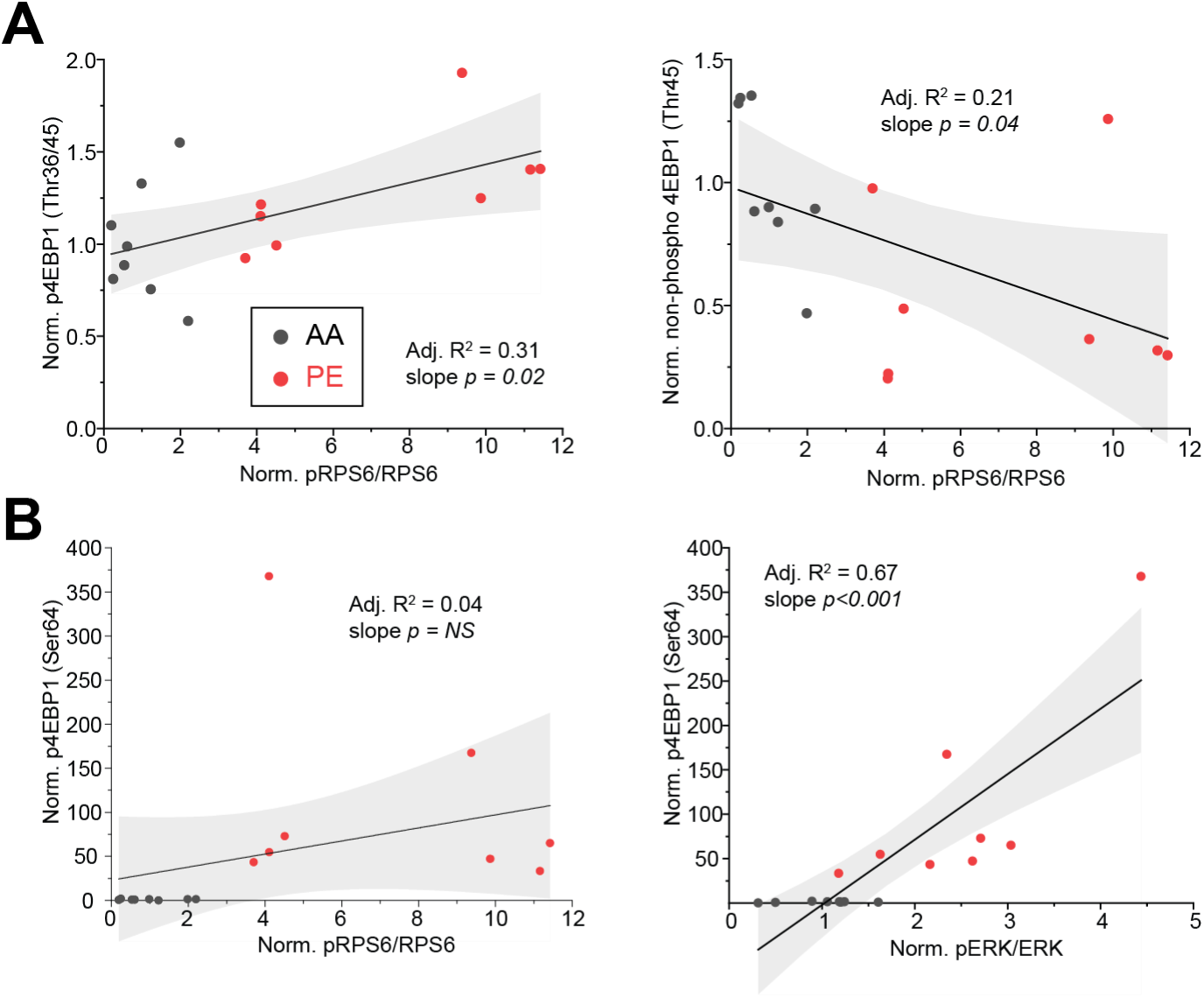
4EP1 phosphorylation at Thr36/45 correlate with mTORC1 activity and 4EBP1 phosphorylation at Ser64 correlates with ERK activity *in vivo*. Heterogeneous activation of mTORC1 and ERK activity were observed following 4 h PE treatment *in vivo*. (A) 4EBP1 phosphorylation at Thr36/45 significantly correlate with mTORC1 activity. Each point represents western blot quantification of the RPS6 phosphorylation and 4EBP1 phosphorylation measured from the same animal. Linear fits of the data are shown in black and the 95% confidence intervals are highlighted in gray. (B) 4EBP1 phosphorylation at Ser64 does not significantly correlate with mTORC1 activity (left) but is strongly correlated with ERK activity *in vivo* (right). Each point represents western blot quantification of the RPS6 phosphorylation and 4EBP1 phosphorylation measured from the same animal. Linear fits of the data are shown in black and the 95% confidence intervals are highlighted in gray.

